# Pan-phylum *In Silico* Analyses of Nematode Endocannabinoid Signalling Systems Highlight Novel Opportunities for Parasite Drug Target Discovery

**DOI:** 10.1101/2022.03.09.483626

**Authors:** Bethany A. Crooks, Darrin McKenzie, Luke C. Cadd, Ciaran J. McCoy, Paul McVeigh, Nikki J. Marks, Aaron G. Maule, Angela Mousley, Louise E. Atkinson

## Abstract

The endocannabinoid signalling (ECS) system is a complex lipid signalling pathway that modulates diverse physiological processes in both vertebrate and invertebrate systems. In nematodes, knowledge of endocannabinoid (EC) biology is derived primarily from the free-living model species *Caenorhabditis elegans,* where ECS has been linked to key aspects of nematode biology. The conservation and complexity of nematode ECS beyond *C. elegans* is largely uncharacterised, undermining the understanding of ECS biology in nematodes including species with key importance to human, veterinary and plant health. In this study we exploited publicly available omics datasets, *in silico* bioinformatics and phylogenetic analyses to examine the presence, conservation and life-stage expression profiles of EC-effectors across phylum Nematoda. Our data demonstrate that: (i) ECS is broadly conserved across phylum Nematoda, including in therapeutically and agriculturally relevant species; (ii) EC-effectors appear to display clade and lifestyle-specific conservation patterns; (iii) filarial species possess a reduced EC-effector complement; (iv) there are key differences between nematode and vertebrate EC-effectors; (v) life stage-, tissue- and sex-specific EC-effector expression profiles suggest a role for ECS in therapeutically relevant parasitic nematodes. These data also highlight putative novel targets for anthelmintic therapies. To our knowledge, this study represents the most comprehensive characterisation of ECS pathways in phylum Nematoda and inform our understanding of nematode ECS complexity. Fundamental knowledge of nematode ECS systems will seed follow-on functional studies in key nematode parasites to underpin novel drug target discovery efforts.

**CONTRIBUTION TO THE FIELD:** This manuscript reports the in silico characterisation of endocannabinoid (EC) signalling pathways across the nematode phylum. The physiological relevance and therapeutic potential of EC signalling in higher organisms has received significant attention. In contrast much of our knowledge on EC signalling in nematodes has been derived from the free-living nematode *Caenorhabditis elegans* where the EC signalling system appears to play key roles in nematode biology and features GPCRs distinct from vertebrate cannabinoid receptors. Unfortunately, the configuration and broader biological significance of EC signalling pathways across the nematode phylum, including in parasites of agricultural, veterinary and medical significance, remains unknown. The *in silico* exploration of the nematode EC signalling system reported here will provide baseline data on novel neuronal signalling pathways to seed future drug target discovery pipelines for parasites.

## INTRODUCTION

Parasitic nematodes inflict a pervasive burden on human, animal and plant health (Robertson et al., 2018). The rapid escalation of anthelmintic resistance, and an over reliance on a limited number of frontline anthelmintics, threatens the global sustainability of parasite control. The need for identification and validation of novel control strategies and chemotherapies for nematode parasites is urgent and requires a robust understanding of unexploited aspects of nematode biology that may offer a source of novel drug target candidates.

Neuromuscular signalling is the primary target for frontline anthelmintics because of its importance to nematode biology. Despite this, many facets of nematode neurobiology, including endocannabinoid signalling (ECS), remain uncharacterised and unexploited for parasite control. The ECS system is a complex lipid signalling pathway involved in the regulation of synaptic transmission via retrograde signalling (Castillo et al., 2012, Zou and Kumar, 2018), and has been associated with a broad range of immunological, psychological, developmental, neuronal and metabolic physiologies in humans where it has significant therapeutic appeal (Kaur et al., 2016). While mammalian (de Azua and Lutz, 2019, Mouslech and Valla, 2009, Fucich et al., 2019) and invertebrate (Acosta-Urquidi and Chase, 1975, Salzet and Stefano, 1998, Stefano and Salzet, 1999, Salzet and Stefano, 2002) ECS pathways have been studied extensively, our knowledge of the presence, structure and function of ECS in nematodes is limited (Aarnio et al., 2011, Galles et al., 2018, Pastuhov et al., 2012, Pastuhov et al., 2016, Batugedara et al., 2018).

In vertebrates, endocannabinoids (ECs) primarily activate the canonical cannabinoid G-protein coupled receptors (GPCRs) CB1 and CB2 (Devane et al., 1992, Mackie, 2008, Sun et al., 2017), in addition to several other cannabinoid-associated receptors (Morales and Reggio, 2017). In contrast, nematodes do not appear to possess homologs of the mammalian-like EC-GPCRs (CB1 and CB2). Instead, the nematode-specific GPCR NPR-19, has been functionally linked to ECS in *C. elegans* (*Ce*-NPR-19) (Oakes et al., 2019, Oakes, 2018, Oakes et al., 2017). *Ce*-NPR-19 displays only 23% sequence similarity with mammalian CB1 but possesses 50% of the key amino acids required for EC ligand (*N*-arachidonoylethanolamine; anandamide; AEA) binding (McPartland and Glass, 2003, Oakes et al., 2017). In addition, *C. elegans* NPR-32 is also activated in response to AEA (Pastuhov et al., 2016), indicating that additional EC-GPCRs could also be present in nematodes.

In vertebrates the primary EC ligands 2-arachidonoylglycerol (2-AG) and AEA are enzymatically biosynthesised on demand, and subsequently metabolised post receptor activation (see Figure 1) (Devane et al., 1992, Ligumsky et al., 1995, Mechoulam and Fride, 1995). 2-AG synthesis occurs via the hydrolysis of diacylglycerol by diacylglycerol lipase (DAGL), while degradation can involve several enzymes including monoacylglycerol lipase (MAGL) and lysophosphatidylserine lipase alpha/beta-hydrolase domain containing-12 (ABHD-12) (see Figure 1A) (Murataeva et al., 2014, Clapper et al., 2018, Blankman et al., 2007). AEA is predominantly synthesised via the hydrolysis of N-arachidonyl phosphatidyl ethanol (NAPE) by N-arachidonyl phosphatidyl ethanol-phospholipase D (NAPE-PLD) and degraded by fatty acid amide hydrolase (FAAH) (see Figure 1B) (Di Marzo, 1999, Biringer, 2021, Liu et al., 2006). There is also evidence to suggest the presence of multiple alternative pathways for EC-ligand biosynthesis and degradation in mammals involving several alternative enzymes including alpha/beta-hydrolase domain containing-4, N-acyl phospholipase B (ABHD-4), lysophospholipase D (Lyso-PLD) and phospholipase A2 (PLA-2) (see Figure 2A) (Liu et al., 2006, Basavarajappa, 2007, Sun et al., 2004).

**Figure 1.**
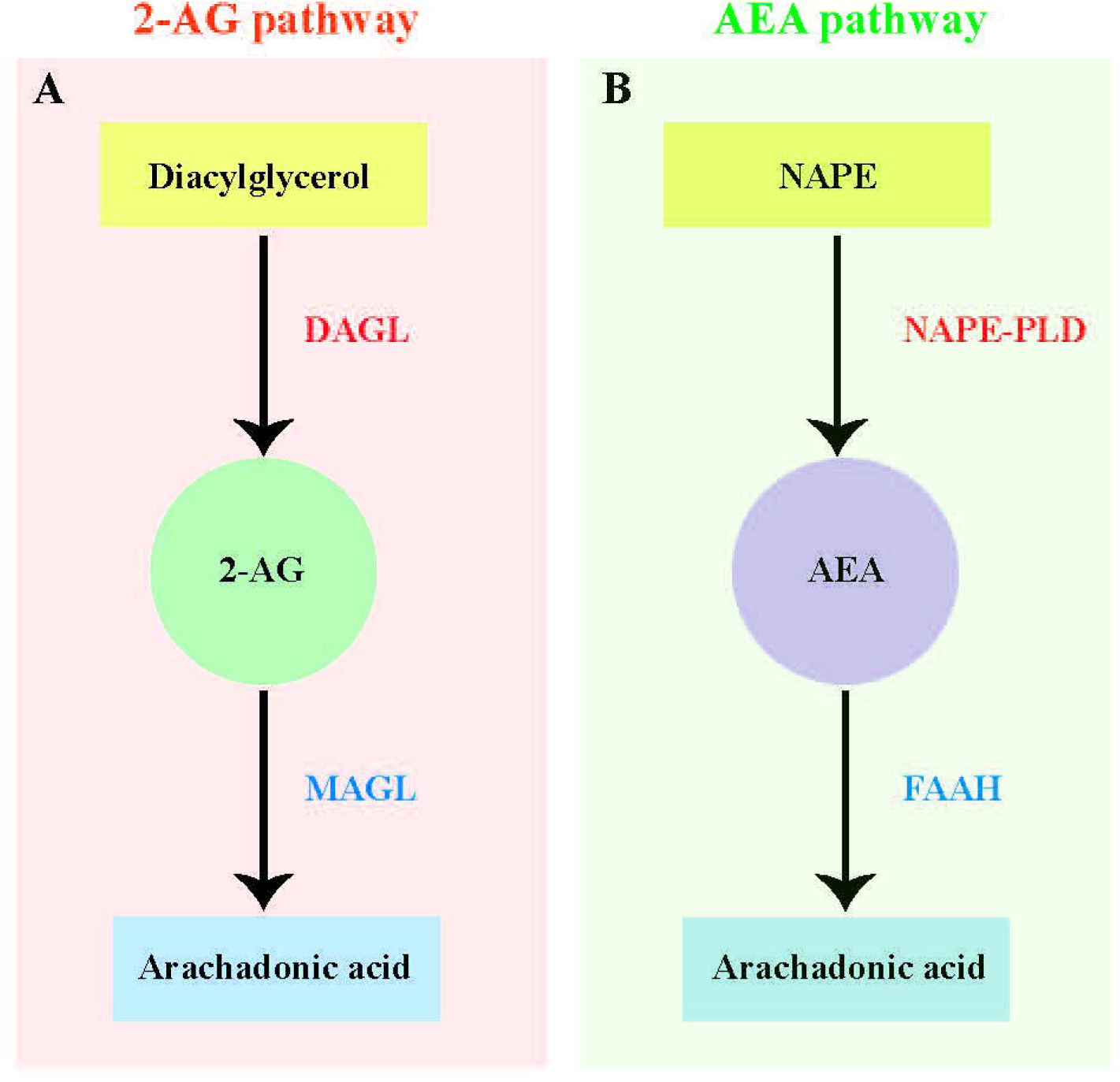
Canonical 2-arachidonoylglycerol and anandamide biosynthesis and degradation pathways. (A) Canonical 2-arachidonoylglycerol (2-AG) biosynthesis and degradation pathway based on vertebrates showing the hydrolysis of diacylglycerol by diacylglycerol lipase (DAGL) and degradation of 2-AG by monoacylglycerol lipase (MAGL). (B) Canonical anandamide (AEA) biosynthesis and degradation pathway based on vertebrates showing the hydrolysis of N-arachidonyl phosphatidyl ethanol (NAPE) by N-arachidonyl phosphatidyl ethanol-phospholipase D (NAPE-PLD) and degradation of AEA by fatty acid amide hydrolase (FAAH).

**Figure 2.**
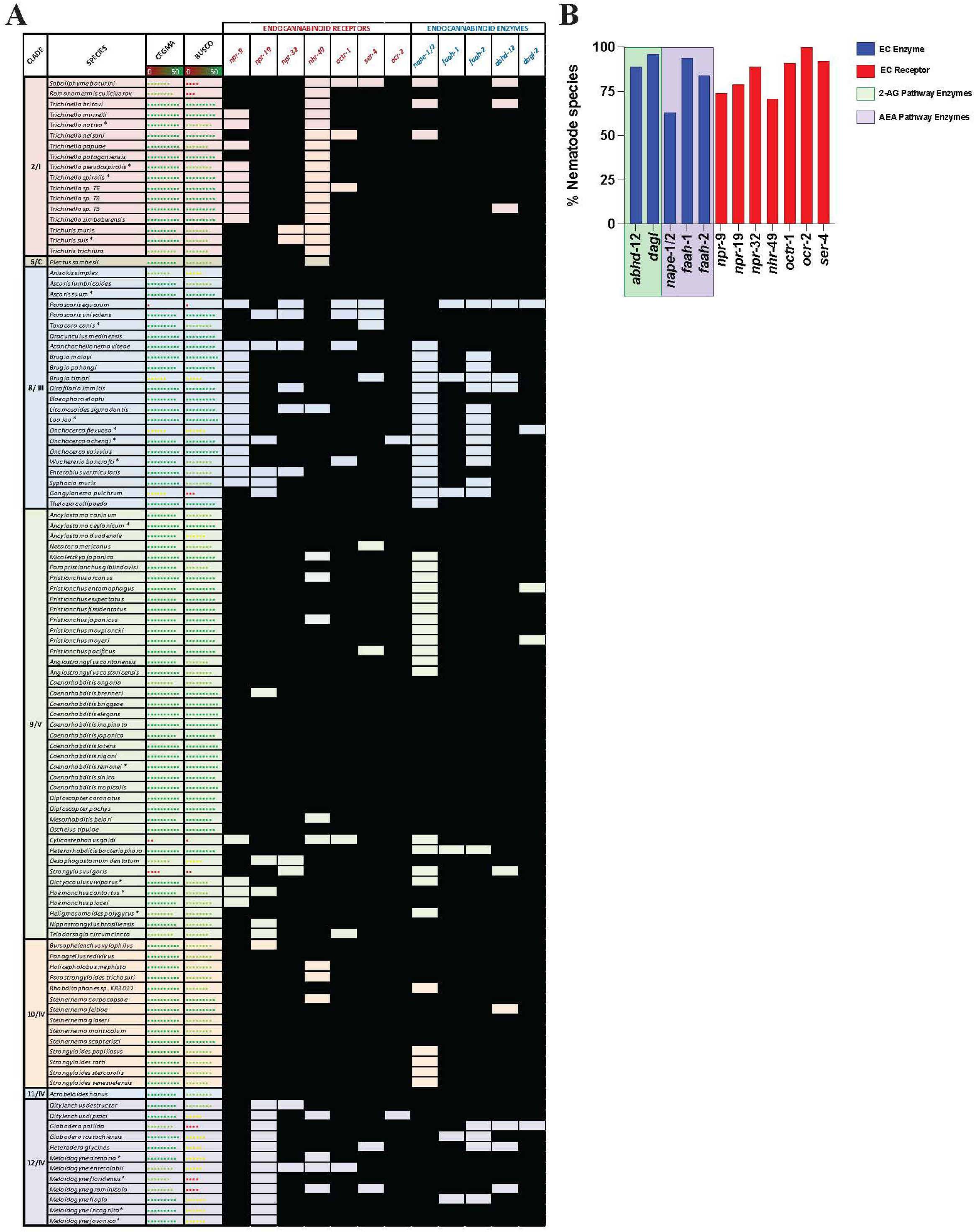
Nematodes possess homologs of canonical endocannabinoid signalling pathway effectors that display broad pan-phylum conservation. (A) Pan-phylum conservation of ECS proteins in all nematodes that possess genome data. Black boxes represent the presence of a homolog. CEGMA/BUSCO scores represent genome quality [data derived from WormbaseParasite, each circle represents 10% increase in genome quality, colours represent scale (red represents lower percentage genome quality, green represents higher percentage genome quality)]. Asterix denotes multiple genomes (*). Nematode genome references listed in File SI 2. All homolog gene IDs identified listed in File SI 3. Clades based on Holterman classification (Holterman et al., 2006). (B) Bar chart displaying the percentage of nematode species that possess each ECR/ECE.

In nematodes 2-AG and AEA have been identified via mass-spectrometry in *C. elegans, Pelodera strongyloides, Caenorhabditis briggsae* and the rodent gastrointestinal nematode *Nippostrongylus brasiliensis* (Lehtonen et al., 2008, Batugedara et al., 2018). In addition, in *C. elegans* several of the hydrolytic enzymes linked to EC degradation (MAGL and FAAH) have been identified *in silico* (Piomelli et al., 2006, Long et al., 2009) and functionally characterised (Lucanic et al., 2011*). Caenorhabditis elegans* ECS has also been associated with a raft of important biological roles (Galles et al., 2018, Lucanic et al., 2011, Pastuhov et al., 2012, Oakes et al., 2019, Oakes et al., 2017).

Data on the presence and function of ECS in parasitic nematodes is limited to a single study that identified ECS enzymes and the putative EC-GPCR NPR-19 via bioinformatics in *N. brasiliensis, Ancylostoma duodenale, Ancylostoma celanicum, Necator americanus, Steinernema carpocapsae, Ascaris lumbricoides, Strongyloides ratti, Strongyloides stercoralis* and *Toxocara canis* (Batugedara et al., 2018). This work also demonstrated that ECs modulate the host immune response during parasite infection and that *N. brasiliensis* produces ECs throughout its lifecycle, most notably in the infective larval stage (Batugedara et al., 2018). This strongly supports the hypothesis that parasitic nematodes possess a functional ECS pathway. However, as these observations represent a small subset (6.7%) of available nematode genomes there remains an opportunity to exploit the recent expansion in nematode omics data to characterise the breadth and complexity of the ECS system across phylum Nematoda.

Here, we employed a bioinformatics driven *in silico* pipeline and phylogenetic analyses to identify the presence, and interrogate the conservation and expression profiles of ECS pathway effectors (EC-effectors) in all publicly available nematodes genomes and life stage and tissue-specific transcriptomes. Our data demonstrate that: (i) ECS is broadly conserved across phylum Nematoda, including in therapeutically and agriculturally relevant species; (ii) EC-effectors appear to display clade and lifestyle-specific conservation patterns; (iii) filarial species possess a reduced EC-effector complement; (iv) sequence analyses reveal key differences between nematode and vertebrate EC-effectors; (v) life stage-, tissue- and sex-specific EC-effector expression profiles suggest a role for ECS in therapeutically relevant parasitic nematodes. To our knowledge this study represents the most comprehensive, pan-phylum, analysis of the nematode ECS system, including in species with global therapeutic and agricultural significance. These data will seed functional genomics studies in tractable parasitic nematodes to inform future novel anthelmintic target discovery pipelines.

## MATERIALS AND METHODS

### Retrieval of query sequences

Query sequences for 70 genes encoding a total of 14 *C. elegans, C. briggsae, Caenorhabditis brenneri, Caenorhabditis japonica* and *Caenorhabditis remanei* endocannabinoid receptors (seven ECRs) and seven endocannabinoid enzymes (seven ECEs) were obtained from WormBase ParaSite v14 (WBP; https://parasite.wormbase.org) (see File SI 1) (Van Gilst et al., 2005, Jose et al., 2007, Harrison et al., 2014, Oakes et al., 2017, Batugedara et al., 2018). Note that in the nematode literature, NAPE-PLD orthologs are commonly referred to as NAPE (McPartland et al., 2006, Lucanic et al., 2011, Harrison et al., 2014) [WormBase Gene IDs; *nape-1* WBGene00021371, *nape-2* WBGene00021370 (Howe et al., 2017)], consequently we have continued to refer to nematode NAPE-PLD as NAPE in this study for consistency.

### Hidden Markov Model and BLAST analysis

A Hidden Markov Model (HMM)-based approach has previously been reported for the identification of flatworm GPCRs (McVeigh et al., 2018). Predicted protein datasets were downloaded from WBP v14 for all nematodes with publicly available genome data (133 genomes; see File SI 2). Predicted protein datasets were concatenated for use as a predicted protein database for HMMERv3.3 HMM-searches. Profile HMMs for nematodes were constructed using predicted protein alignments of all candidate EC-effector protein homologs in *C. elegans, C. briggsae, C. brenneri, C. japonica* and *C. remanei* (see File SI 1). Multiple Sequence Alignments (MSAs) were generated using EMBL-EBI Clustal-Omega [https://www.ebi.ac.uk/Tools/msa/clustalo; (Sievers et al., 2011)]. The *hmmsearch* function was employed to identify putative EC proteins within the nematode predicted protein datasets (see File SI 2) using default settings. Due to the volume of genomic data generated, the highest confidence hits were selected for each protein based on an inclusion threshold of E-value ≤0.01 and/or a score of ≥150.

Putative EC protein sequences identified via *hmmsearch* were then used as queries in reciprocal BLASTp searches of WBP (https://parasite.wormbase.org/Multi/Tools/Blast; default settings) and NCBI non-redundant (https://blast.ncbi.nlm.nih.gov; default settings) databases. Queries that failed to return a putative ECR or ECE hit within the top 4 BLAST results were excluded from further downstream analyses. All BLASTp searches returning negative hits (hits outside of the outlined inclusion criteria and those that returned no hits) were confirmed negative via tBLASTn searches; this approach also mitigated the impact of poor genome quality (false negatives) on our analyses where relevant.

### Post-BLAST sequence analysis

Key EC ligand binding residues and functional motifs for mammalian and invertebrate ECRs and ECEs were identified from the published literature (see File SI 3) and all positive hits were examined visually for the presence of any key residues/motifs via multiple sequence alignments using EMBL-EBI Clustal-Omega [https://www.ebi.ac.uk/Tools/msa/clustalo (Sievers et al., 2011)]. The presence of known protein family or structural domains in ECE BLAST hits was analysed using InterProScan [https:www.ebi.ac.uk/interpro/search/sequence-search, (Jones et al., 2014)]. Putative ECR hits were analysed for the presence of GPCR transmembrane domains using EMB TMpred server [https://embnet.vital-it.ch/software/TMPRED_form.html (Hofmann, 1993)]. Any putative ECE hits which lacked the required family/protein domains for designation as an ECE, or putative ECR sequences which possessed ≤3 transmembrane domains, were excluded from further analysis.. ECRs and ECEs were analysed for the presence of conservative substitutions of key residues or within binding motifs using WebLogo3 (Schneider and Stephens, 1990, Crooks et al., 2004) (see Table SI 2).

### Phylogenetic analysis

MUSCLE was used to create multiple sequence alignments (MSAs) of protein sequences for all positive EC protein hits using MEGA X (Stecher et al., 2020). For ECRs, alignments were manually edited to include only TM domains, for ECEs only functional domains were included in analysis. Functional domains for ECEs and transmembrane domains for ECRs were identified via the NCBI Conserved Domains Database [CDD; https://www.ncbi.nlm.nih.gov/Structure/cdd/wrpsb.cgi (Lu et al., 2020b)]. Maximum likelihood (ML) phylogenetic trees were constructed using PhyML [http://www.phylogeny.fr (Dereeper et al., 2008)] from the domain only MUSCLE MSAs with default parameters and branch support assessment using the approximate likelihood ratio test (aLRT) with “SH-like” parameters. Trees were exported from PhyML in Newick format and were drawn and annotated using the Interactive Tree of Life [iTOL; https://itol.embl.de (Letunic and Bork, 2021).

### Transcriptome analysis

180 publicly available transcriptome datasets [145 life stage- and 35 tissue-specific datasets] representing 32 nematode species were analysed in this study. One hundred and fifty publicly available life stage and tissue specific transcriptome datasets representing 27 nematode species were collated from WBP v14 Gene Expression database (Howe et al., 2017) and published literature (see File SI 2). WBP datasets (see File SI 2) consisted of metadata, raw counts, transcripts per million (TPM) and DESeq2 differential expression data (in log2foldchange and adjusted p value formats). Data for an additional 34 datasets, representing five species [*Haemonchus contortus, Toxocara canis, Globodera pallida, Strongyloides venezuelensis* and *Strongylodies papillosus*; see File SI 2] were identified from published literature, and metadata and raw counts were accessed from NCBI Sequence Read Archive [SRA; www.ncbi.nlm.nih.gov/sra] (National Centre for Biotechnology Information, 2009) for analysis. TPM and median TPM data for *H. contortus, T. canis* and *G. pallida* were downloaded using the European Bioinformatics Institute RNAseq-er Application Program Interface (Petryszak et al., 2017) (File SI 2). *S. venezuelensis*and *S. papillosus* (raw counts and TPM) data were analysed using an established RNA-Seq pipeline (Lu et al., 2020a). Briefly, raw sequences reads were processed into forward and reverse fastq files using the NCBI SRA Toolkit (SRA Toolkit Development Team, 2020). Reads were then trimmed using Trimmomatic (v0.36; parameters: LEADING:5 TRAILING:5 SLIDINGWINDOW:3:15 MINLEN:34) (Bolger et al., 2014) and sequences below this established inclusion threshold were removed. Corresponding genome assemblies [BioProject accessions; PRJEB530 and PRJEB525, respectively] (Hunt et al., 2016) were downloaded from WormBase ParaSite v14 (Howe et al., 2017) and reads were mapped to the relevant genome using HISAT2 v2.1.0 (Kim et al., 2015). Raw gene counts were assigned via SubRead v 2.0.1 featureCounts (Liao et al., 2014). Raw counts of orthologous genes were transformed to TPM and subsequently median TPMs were calculated to represent raw gene expression in the life stages with RNA-Seq data available. An inclusion threshold for expression of 1.5 TPM was applied [TPM thresholds are typically set between 1-2 TPM (Wagner et al., 2013, Soneson and Robinson, 2018)], any transcripts which failed to meet the threshold for expression were excluded from downstream analysis.

Differential expression data for *H. contortus, T. canis, G. pallida, S. venezuelensis* and *S. papillosus* RNA-Seq data were generated using DESeq2 in the format of log2foldchange and adjusted p values (Love et al., 2014, RS Team, 2015). Datasets were then mined for pathway protein gene IDs identified in the HMM searches. Heatmaps displaying log2AverageTPM were generated using the Heatmapper Expression protocol, with an average linkage clustering method and Pearson’s distance measurement method (Babicki et al., 2016).

## RESULTS AND DISCUSSION

### Nematodes possess homologs of canonical endocannabinoid signalling pathway effectors

In this study we mined 133 nematode genomes (representing 109 species, 7 clades and 3 lifestyles) for 13 putative ECS pathway effectors, expanding upon previous studies (Batugedara et al., 2018). Our HMM-based *in silico* approach returned a total of 1289 putative ECS effector homologs (ECEs and ECRs) (see Figure 2 and File SI 3). The data demonstrate that: (i) ECEs and ECRs display pan-phylum conservation; (ii) representatives of Clades 8 and 12 exhibit the lowest level of EC-effector conservation [clade 8: 68% and 79% of all possible ECEs and ECRs conserved, respectively; clade 12: 83% and 72% of all possible ECEs and ECRs conserved, respectively] and, (iii) free-living and parasitic nematodes display a comparable level of EC-effector conservation [free-living: 91% and 96% of all possible ECEs and ECRs conserved, respectively; parasitic: 83% and 92% of all possible ECEs and ECRs conserved, respectively] (see Figure 2). To our knowledge this is the first pan-phylum examination of nematode ECS profiles and represents a comprehensive analysis of ECS pathway conservation in parasitic species that impact human, animal and plant health. Several important points emerge from these data (see below).

### Endocannabinoid signalling system effectors are broadly conserved across phylum Nematoda

Nematode EC-effectors (ECEs responsible for synthesis and degradation of EC ligands, and putative ECRs) appear to be broadly conserved across phylum Nematoda (Figure 2A). ECEs display greater conservation than putative ECRs across all nematodes, with the exception of the ECE *nape*-1/2 which appears to be absent from 37% of nematodes, many of which are representatives of clades 8 and 9 (Figure 2A and B). The more conserved profile of ECEs versus ECRs is consistent with the requirement for specific ECEs in the biosynthesis and degradation of EC-ligands and the potential for redundancy among ECRs, which has been documented in other systems (Paulsen and Burrell, 2019, Lu and Mackie, 2021).

Genes encoding the key ECEs responsible for 2-AG synthesis and metabolism, DAGL-2 and ABHD-12 respectively, are co-conserved in 87% of nematodes (see Figure 2), indicating that a significant proportion of nematode species are likely to possess a functional, canonical, 2-AG synthesis and degradation pathway. Most (95%) nematode species examined possess an gene ortholog of the 2-AG biosynthesis enzyme DAGL-2, while the gene encoding ABHD-12, responsible for catalysing 2-AG metabolism is present in 88% of species. *Parascaris equorum* and *Globodera pallida* do not appear to possess either a *dagl-*2 or *abhd-*12 homolog, however their genome quality is lower as indicated by CEGMA/BUSCO scores [see Figure 2; (Parra et al., 2009, Simão et al., 2015)].

Genes encoding NAPE and FAAH, the primary enzymes responsible for the synthesis and metabolism of AEA, are co-conserved in 55% of nematode species, suggesting that a significant proportion nematodes have the ability to synthesise and degrade AEA (Figure 2). This is corroborated by studies that have identified AEA in several nematodes, including *C. elegans* and *N. brasiliensis*, via mass-spectrometry (Lehtonen et al., 2008, Batugedara et al., 2018). While 88% of species encode at least one FAAH homolog, the NAPE-encoding gene is conserved in only 63% of nematodes (either *nape*-1 or -2). Notably, many of the species that lack a *nape* homolog are filarial nematodes, *Strongyloides* species or members of the Diplogasteroidea superfamily (Figure 2). Whilst the absence of *nape* in some species may be explained by genome quality, in other species with robust genome data an alternative AEA synthesis pathway may exist (see below). Previous studies have demonstrated the presence of two, functionally divergent, *nape* orthologs in *C. elegans* that occupy adjacent genomic positions (*nape*-1, <u>IV:3739520..3740880</u>; *nape*-2, IV:3735470..3738925; [WormBase; (Davis et al., 2022)], share 73% sequence identity, and display complete conservation of the NAPE-PLD signature sequence (Harrison et al., 2014). Our pan-phylum analysis confirms that *C. elegans* is the only nematode species that possesses two distinct NAPE-encoding genes. In all other species that encode NAPE the same gene ID was returned for both the *nape*-1 and *nape*-2 BLASTp searches (one positive *nape* hit was considered a positive return for both *nape*-1 and *nape*-2 and was designated *nape*-1/2; see Figure 2 and File SI 3). The presence of a single *nape-*1/2 in all of the nematodes examined, including other *Caenorhabditis* species, suggests that *nape*-1 and -2 may have arisen as a result of a relatively recent gene duplication event in *C. elegans* (Harrison et al., 2014).

Of the seven putative ECRs included in this study NPR-19 and NPR-32 have been most closely linked to ECS in nematodes (Pastuhov et al., 2012, Pastuhov et al., 2016, Oakes et al., 2017). Our *in silico* analysis reveals that 79% of the nematode species investigated in this study possess NPR-19 and 89% possess NPR-32, underscoring their putative importance to nematode biology. Beyond NPR-19 and -32, OCR-2 an ortholog of the human transient receptor potential vanilloid channel (TRPV) (de Bono et al., 2002), has also been closely linked to ECS. OCR-2 regulates signal transmission and thus modulates several *C. elegans* behaviours (Lee and Ashrafi, 2008, Oakes et al., 2019). It is interesting to note that 100% of nematode species examined in this study possess a gene encoding OCR-2, also suggesting a significant role in nematode biology. NPR-9 has been implicated in locomotion, regulation of innate immune responses, roaming and foraging behaviours in *C. elegans*(Bendena et al., 2008, Campbell et al., 2016, Yu et al., 2018), while GPR-55, the human ortholog of nematode NPR-9, is known to interact with human CB1 and CB2 to form functionally important heteromers (Balenga et al., 2014, Kargl et al., 2012). The lower level of *npr*-9 conservation across phylum Nematoda revealed here (*npr*-9 conserved in 60% of species examined; Figure 2) may indicate a less conserved functional role for this putative ECR in nematodes.

### Nematode EC-effector conservation profiles display clade specific trends

Our data demonstrate distinct conservation patterns of EC-effectors across nematode clades. Whilst clade 9 and 10 nematodes exhibit the highest degree of EC-effector conservation, clade 8 species display the most reduced complements (Figure 2). This could be, in part, explained by lower CEGMA/BUSCO scores for clade 8 genomes, such that the profiles presented here may not be a true representation of EC-effector complements in this clade. Variable genome quality is an inherent caveat to *in silico* approaches such as those employed in this study however, the use of tBLASTn for all negative BLASTp returns can help to mitigate this, in addition to continued improvements in genome quality (Doyle and Cotton, 2019).

For other clades there appear to be ECR specific trends. For example, *npr-*32 is broadly conserved across all clades, pointing towards a more conserved function for this receptor pan-phylum. Other ECRs display more clade-specific, restricted profiles, including *nhr*-49 which appears to be entirely absent from clade 2 (Figure 2A). Clade 2 nematode genome assemblies are high quality (as indicated by CEGMA/BUSCO scores; (Howe et al., 2017) and likely provide a true reflection of the *nhr*-49 profile. Notably, *npr-19* is also entirely absent from clade 12 species, indicating that alternative ECRs may contribute to the ECS pathway in these nematodes. Indeed, clade 12 nematodes exhibit broad conservation of other putative ECRs (e.g. *npr-*9, -32, *nhr-*49, *octr-*1, *ser-*4, *ocr-*2; see Figure 2A).

### Filarids possess distinctive EC-effector profiles

Whilst there appear to be limited differences between free-living and parasitic nematode EC-effector profiles in general [96% ECR conservation in of free-living nematodes (FL) vs 92% in parasitic species; 91% ECE conservation in free-living nematodes vs 83% in parasitic species], filarial nematodes display distinct EC-effector gaps. For example, filarial species completely lack genes encoding NPR-9 and NAPE-1/2, and have a significantly reduced FAAH-2 encoding gene profile (Figure 2A). The absence of NAPE-1/2 points towards the presence of an alternative AEA synthesis pathway in filarids, which may also be the case for other species e.g. *Pristionchus* spp. and *Strongyloides* spp. that likewise lack NAPE-1/2 encoding genes (see below; Figure 2A).

### Nematodes that lack NAPE-1/2 possess putative alternative AEA synthesis enzymes

Our data indicate that 41 nematode species lack genes encoding the nematode NAPE-PLD ortholog NAPE-1/2, the enzyme primarily responsible for AEA synthesis (Figure 2A) (Di Marzo, 1999). In vertebrates, two additional pathways have been implicated in the synthesis of AEA: (i) hydrolysis of NAPE by ABHD-4 forming the intermediate glycerophosphoanandamide (Glycero-p-AEA) which, following further hydrolysis by glycerophosphodiester phosphodiesterase 1 (GDE-1), results in AEA (see Figure 3A ii); (ii) hydrolysis of NAPE by phospholipase A2 (PLA-2) to form the intermediate N-acyl-1-acyl-lyso-PE (lyso-NAPE) and subsequently, following hydrolysis by lysophospholipase D (Lyso-PLD), results in AEA (see Figure 3A iii) (Wang and Ueda, 2009, Muccioli, 2010, Simon and Cravatt, 2006, Astarita and Piomelli, 2009). To determine if alternative AEA synthesis pathways exist in nematodes that appear to lack the classical NAPE-1/2 AEA biosynthesis pathway, we mined all available genome data for *abhd-*4, *gde-*1, *pla-*2 and *lyso-PLD*.

**Figure 3.**
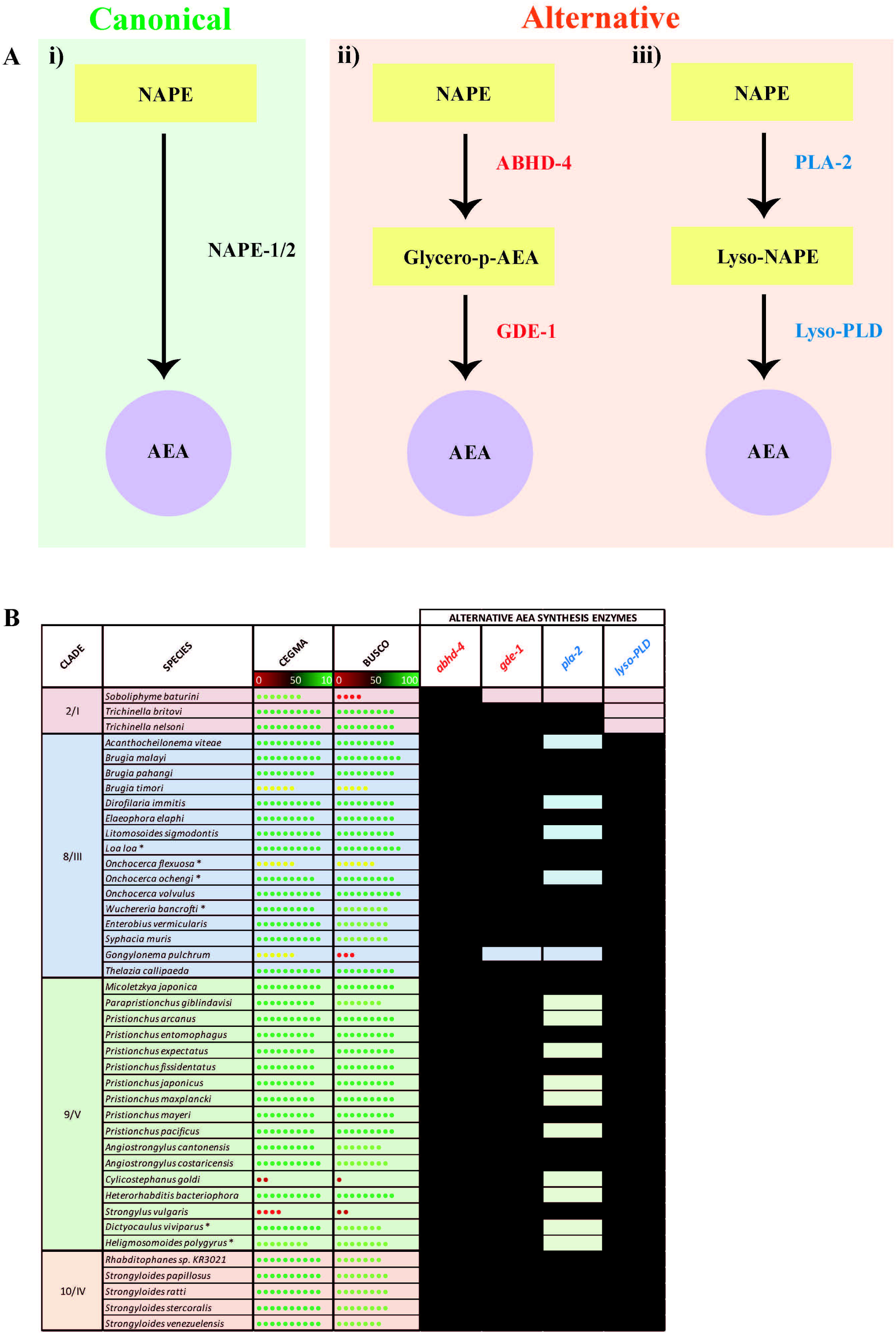
Nematode species lacking the anandamide synthesis enzyme NAPE possess putative alternative AEA synthesis enzymes. (A) Diagram showing canonical (i) anandamide (AEA) synthesis pathway alongside two alternative synthesis pathways: (ii) hydrolysis of NAPE by ABHD-4 forming the intermediate glycerophosphoanandamide (Glycero-p-AEA) and hydrolysis by glycerophosphodiester phosphodiesterase 1 (GDE-1) to synthesise AEA, and (ii) hydrolysis of NAPE by phospholipase A2 (PLA-2) to form the intermediate N-acyl-1-acyl-lyso-PE (lyso-NAPE) and hydrolysis by lysophospholipase D (Lyso-PLD) to synthesise AEA. (B) Conservation of genes encoding the alternative AEA synthesis enzymes ABHD-4, GDE-1, PLA-2, Lyso-PLD is shown in all nematodes that lack NAPE. Black boxes represent the presence of a homolog. CEGMA/BUSCO scores represent genome quality [data derived from WormbaseParasite, each circle represents 10% increase in genome quality, colours represent scale (red represents lower percentage genome quality, green represents higher percentage genome quality)]. Asterix denotes multiple genomes (*). Nematode genome references listed in File SI 2. All homolog gene IDs identified listed in File SI 3. Clades based on Holterman classification (Holterman et al., 2006).

Our data reveal that 39 of the 41 species that lack NAPE-1/2 possess at least one putative alternative AEA synthesis pathway (see Figure 3B, File SI 3). 95% of these species possess both *abhd*-4 and *gde-*1 (alternative pathway shown in Figure 3A ii), while 56% encode both PLA-2 and Lyso-PLD (alternative pathway shown in Figure 3A iii); 56% of nematodes encode the enzymes for both alternative AEA synthesis pathways. Therefore these data suggest that nematodes which lack the classical NAPE-1/2 biosynthesis pathway predominantly synthesise AEA via ABHD-4 and GDE-1. However, within clades 8, 9 and 10 there are examples of species which may have the ability to synthesise AEA via either alternative pathway e.g. *Strongyloides* species and several filarial nematodes (Figure 3B).

Interestingly in mammals, in addition to PLA-2, other enzymes have been linked to the synthesis of lyso-NAPE (Sun et al., 2004), for example, ABHD-4 can remove an acyl group from NAPE resulting in the creation of lyso-NAPE (Simon and Cravatt, 2006). Therefore the presence of *abhd-4* in 100% of the nematodes investigated in this study suggests that some nematodes that possess *lyso-PLD* but lack *pla*-2 could compensate by employing ABHD-4 in the synthesis of lyso-NAPE.

*In silico* evidence for the presence of putative, alternative, AEA biosynthesis enzymes in a range of therapeutically relevant nematodes strongly suggests that NAPE-1/2 independent pathways contribute to AEA synthesis in these species. Further analysis, including mass spectrometry to isolate AEA, will begin to unravel the importance of alternative AEA biosynthesis pathways in these nematodes.

### Nematode FAAH homologs display conservative substitutions in a key AEA binding site

The mammalian AEA hydrolysis enzymes FAAH-1 and FAAH-2 possess four key residues required for catabolic activity (FAAH-1: K142, M191, S217, S241; FAAH-2: K131, C180, S206, S241) (Lucanic et al., 2011, Sirrs et al., 2015, Haq and Kilaru, 2020). Analysis of nematode FAAH-1 homologs identified in this study (see Figure 2) revealed that >90% possess the key mammalian binding site residues K142, S217 and S241, while 83% of identified nematode FAAH-2 homologs possess K131, S206 and S241 (see Figure 4A-C). However, 87% of nematode FAAH-1 homologs display a conservative substitution (methionine for leucine) at position 191 while 80% of nematode FAAH-2 homologs substitute cysteine for leucine at position 180 (Figure 4A-C). M191 has been implicated in mammalian FAAH-1 EC-derivative binding, with studies demonstrating lipophilic interactions between this residue and partial cannabinoid receptor agonists, linking it to AEA binding (Chicca et al., 2018), however the significance of C180 (FAAH-2) is less clear (Janowitz et al., 2003, Rossignoli et al., 2018, Facchiano and Marabotti, 2010).

**Figure 4.**
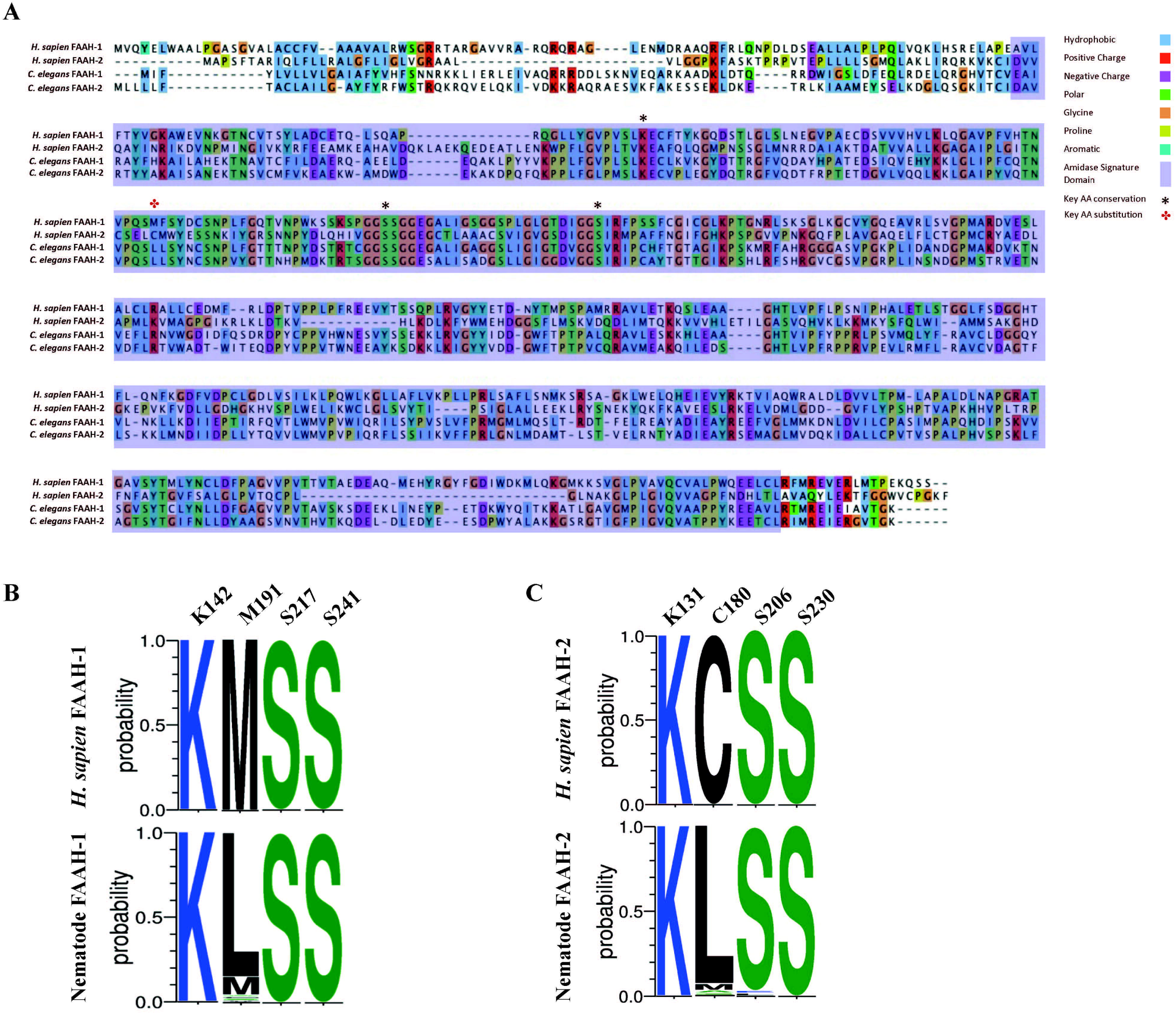
Nematode FAAH homologs conserve key functional domains and motifs, but display conservative substitutions in a key AEA binding site. (A) Protein sequence alignment of *Homo sapien* FAAH-1, *H. sapien* FAAH-2, *Caenorhabditis elegans* FAAH-1 and *C. elegans* FAAH-2. Amino acids are highlighted in the same colour if > 60% of residues are conserved. Legend denotes conserved amino residues, key ligand binding residues in FAAH-1 and FAAH-2, and the amidase signature domain. AA denotes amino acid; *H. sapien* denotes *Homo sapien*, *C. elegans* denotes *Caenorhabditis elegans*. (B) Amino acid sequence-logo representing sequence diversity between key residues in nematode FAAH-1 homologs (consensus) vs *H. sapien* FAAH-1 [O00519, FAAH1_HUMAN] and, (C) Amino acid sequence-logo representing sequence diversity between key residues in nematode FAAH-2 homologs vs *H. sapien* FAAH-2. Key EC binding site residues were derived from *H. sapien* FAAH-1 and -2 and are detailed in the top column of each table. Colours indicate hydrophobicity of amino acid residues (hydrophilic residues are blue, neutral residues are green and hydrophobic residues are black).

Mammalian FAAH-1 and -2 also possess a highly conserved 130 bp amidase signature domain that enables enzyme characterisation (Ahn et al., 2009, Chebrou et al., 1996); this domain is conserved in 94% of the nematode species examined in this study (Figure 4A).

While these data demonstrate that nematodes possess homologs for FAAH enzymes that display broad conservation with vertebrates, the presence of a distinct substitution in a key AEA binding site across many nematodes may highlight the potential for drug target selectivity towards parasitic nematode species.

### Nematodes possess homologs for the mammalian 2-AG degradation enzyme ABHD-12

In mammals, MAGL is primarily responsible for 2-AG degradation and thus the termination of EC signalling (Murataeva et al., 2014, Kano et al., 2009). Additional 2-AG degradation enzymes, ABHD-5 -6 and -12, have also been reported (Savinainen et al., 2012, Blankman et al., 2007) however, ABHD-5 and -6 are thought to have a less significant role in 2-AG degradation (Murataeva et al., 2014, Kano et al., 2009).

The nematode literature presents conflicting data on the identity of the 2-AG degradation enzyme; indeed prior to this study it was unclear whether nematodes possess a true ortholog of MAGL (Oakes et al., 2019, Oakes et al., 2017) or, if the nematode 2-AG degradation enzyme is actually an ABHD-12 ortholog (Batugedara et al., 2018, Clarke et al., 2021). It is interesting to note that previous work indicates that *magl* homologs are absent in some nematode species (Batugedara et al., 2018), which has been confirmed here using our pan-phylum *in silico* approach (Figure 2A). Indeed, in our analyses only two nematode species (of the 109 in this study) returned an *magl*-like sequence within the top 5 BLASTp hits (*Strongylus vulgaris,* SVUK_0001964001; *Steinernema scapterisci*, L892_g30127.t1), both of which failed to meet E-value inclusion criteria. The absence of MAGL in nematodes indicates the presence of an alternative 2-AG degradation enzyme.

In light of the limited role for ABHD-5 and -6 in mammalian 2-AG degradation and the absence of MAGL across nematodes (as reported here and in previous studies), we focused our attention on ABHD-12 as a putative alternative to MAGL in nematodes. Pan-phylum analysis of nematode genomes identified orthologs for ABHD-12 in 88% of nematode species examined. Significantly, phylogenetic analyses of these putative *abhd-12* homologs demonstrated that 99% of nematode BLAST returns for *abhd-12* cluster more strongly with human *abdh-12* than human *magl* (Figure 5 and Figure SI 1) suggesting that the nematode 2-AG hydroxylase enzymes identified here, and originally designated as *magl* in *C. elegans* [Y97E10AL.2; (Oakes et al., 2019)], are orthologous with *abhd-12*.

**Figure 5.**
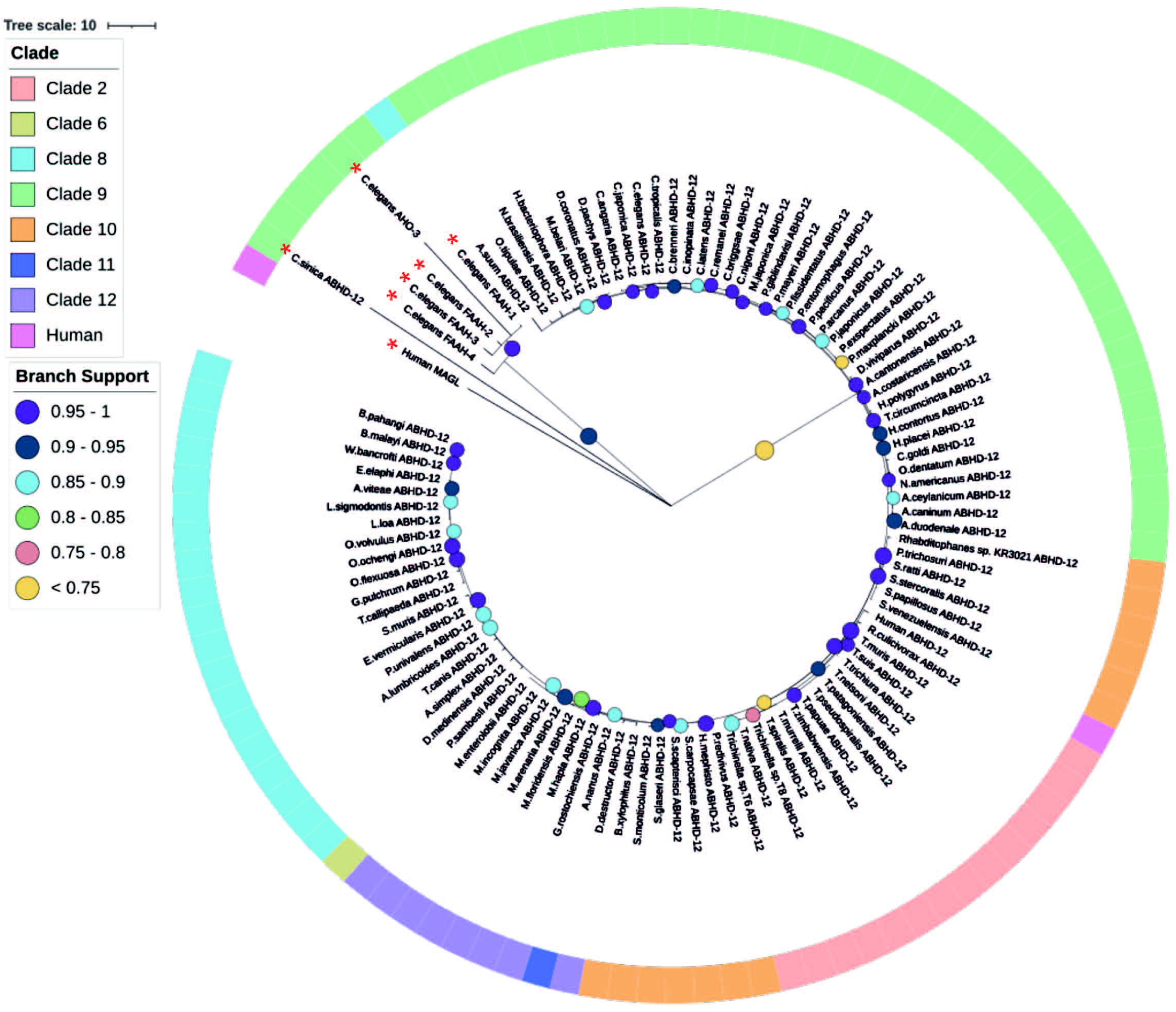
Maximum likelihood phylogeny of nematode ABHD-12 homologs. 98 nematode ABHD-12 homologs are shown in addition to *Homo sapien* ABHD-12 [Q8N2K0 (ABD12_HUMAN)] *H. sapien* MAGL [Q99685 (MGLL_HUMAN)], *Caenorhabditis elegans* FAAH-*1-4* [WBGene00015047, WBGene00015048, WBGene00019068, WBGene00013232] and *C. elegans* AHO-3 [WBGene00045192; alpha/beta hydrolase containing protein ]. Non-ABHD-12 homologs are marked with a red asterisk (*). Outer ring denotes nematode clade and coloured circles represent branch support values. Tree was generated from an alignment trimmed to include protein functional domains. Branch supports indicate statistical support from approximate likelihood ratio test (aLRT).

### Nematode NPR-19 and -32 orthologs possess key functional motifs and EC binding residues

NPR-19 has been identified as a putative EC-GPCR in *C. elegans* and appears to modulate key aspects of nematode biology including axon regeneration, locomotion, modulation of noiceception and feeding, and therefore may represent a promising anthelmintic target (Oakes et al., 2017).

78% of the nematodes examined in this study possess an *npr-19* ortholog (see Figure 2 and Figure 6A). Phylogenetic analysis demonstrates that nematode *npr-19* orthologs failed to cluster with human CB1 and CB2, confirming that NPR-19 is not the direct ortholog of the human EC-GPCRs receptors (CB1 and CB2 ) (Figure 6A and Figure SI 1). Further analysis of the nematode *npr-19* orthologs revealed that whilst the nematode NPR-19 consensus sequence has only 23% similarity to human CB1, several of the known human CB1 EC binding residues (N46, D88, F189, L193, F379, S383) (McAllister et al., 2003, Reggio, 2010) are conserved (Figure 6B). This indicates that nematode NPR-19 orthologs are likely to possess EC ligand binding capacity and aligns with previous work in *C. elegans* (Oakes et al., 2017). When examined at the clade level, sequence analysis showed that two conservative substitutions are present at position K192 (82% of clade 2 species substitute K192 for N192), while species in clades 8, 9, 10 and 12 substitute K192 for D192. K192 forms a hydrogen bond with the amide oxygen of AEA, implicating this residue in ligand binding (McAllister et al., 2003) (see Figure 6B).

**Figure 6.**
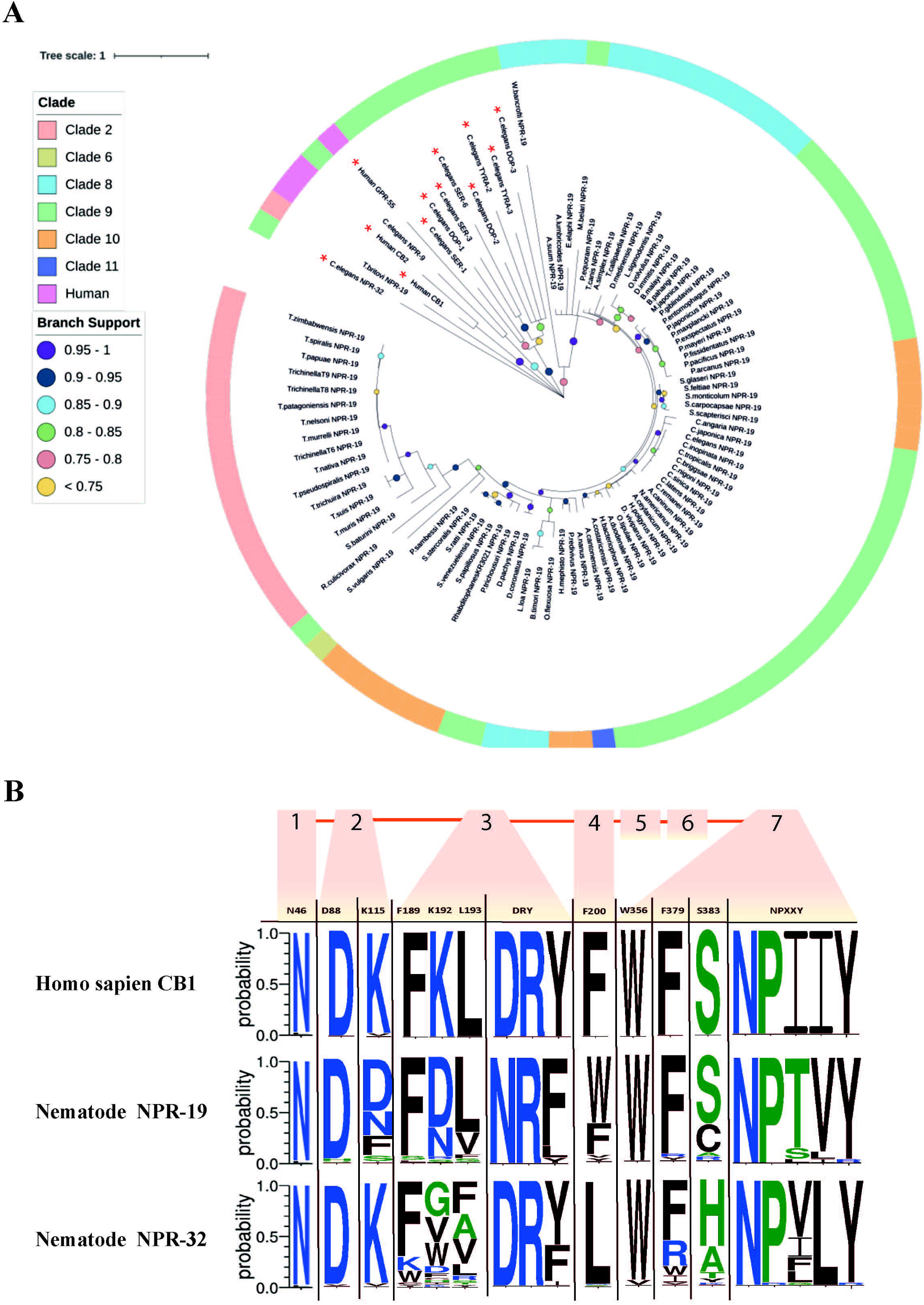
Maximum likelihood phylogeny of nematode *npr-19* homologs. 85 nematode NPR*-*19 homologs are shown in addition to *Homo sapien* CB1 and CB2 [P21554 (CNR1_HUMAN), P34972 (CNR2_HUMAN)], *H. sapien* GPR-55 [Q9Y2T6 (GPR55_HUMAN)] and several *Caenorhabditis elegans* biogenic amine receptors (serotonin [SER-1,-3,-6; WBGene00004776, WBGene00004778, WBGene00021897] dopamine [DOP-1-3; WBGene00001052, WBGene00001053, WBGene00020506] and tyramine [TYRA-2 and -3; WBGene00017157, WBGene00006475]). Non-NPR*-*19 homologs are marked with a red asterisk (*). Outer ring denotes nematode clade and coloured circles represent branch support values. Tree was generated from an alignment trimmed to include functional domains. Branch supports indicate statistical support from approximate likelihood ratio test (aLRT). (B) Amino acid sequence-logo demonstrating sequence diversity between nematode NPR-19 and NPR-32 orthologs (consensus) and *H. sapien* CB1 [P21554 (CNR1_HUMAN)]. Known vertebrate endocannabinoid binding and GPCR motifs/residues are indicated in the top row of the sequence logo table, transmembrane regions 1-7 are indicated by orange boxes and numbers, amino acid colours indicate hydrophobicity of amino acid residues (hydrophilic residues are blue, neutral residues are green and hydrophobic residues are black).

NPR-32 is implicated in *C. elegans* axon regeneration (Pastuhov et al., 2016) and, in addition to NPR-19, is believed to be a putative EC-GPCR (Oakes et al., 2017). In this study we identified 97 NPR-32 homologs (Figure 2 and Figure SI 1) which share (consensus sequence) only 21% identity with the human EC-GPCR CB1, but conserve several key residues (N46, D88, K115, F189; Figure 6C) that are believed to be important for ECS function (Pastuhov et al., 2016).

In addition, CB1 possesses a “toggle switch” (residue W356), a putative molecular hinge that interacts with F200 to change the form and state of the receptor which in turn aids EC ligand binding (Pei et al., 2008, Al-Zoubi et al., 2019). In nematodes the “toggle switch” W356 is conserved in >85% of NPR-19 and -32 orthologs, whereas NPR-19 F200 is substituted for W200 in 59% of species and NPR-32 F200 is substituted for L200 in 90% of species, (see Figure 6C). The significance of these observations will be revealed through molecular docking studies, crystal structure analysis, and functional genomics in relevant parasite species, and will inform the role and importance of these receptors in nematode ECS biology.

### EC-effectors are differentially expressed across nematode life stages, sexes and tissues, suggesting key roles in parasite biology

EC-effector expression has previously been examined in several parasites demonstrating differential expression across life-stages (Batugedara et al., 2018). Here we further profiled EC-effector expression in 32 nematode species representing several distinct lifestyles (see File SI 2).

Our data demonstrate that several of the putative ECRs examined in this study are upregulated in third-stage larvae (L3) of several parasites including *Ancylostoma ceylanicum, Teladorsagia circumcincta, Dictyocaulus viviparus, H. contortus, S. ratti, S. stercoralis* and *Onchocerca volvulus* (see Figure SI 2). Parasitic nematode L3 larvae are analogous to the dauer life stage of *C. elegans*; both *C. elegans* dauer and parasitic nematode L3 stages display similar physiology, are in arrested development, are non-feeding and are highly resistant to their environment (Hotez et al., 1993, Viney et al., 2005, Crook, 2014). L3 parasites of species such as those outlined above transition from arrested (dauer-like) L3 larvae to infective L3 (iL3) larvae either constitutively or via the influence of host and environmental factors (REF). The ECS pathway has been implicated in antagonization of dauer formation and abolishing dauer larval arrest via stimulation of cholesterol in *C. elegans* and, in turn, promotion of nematode growth and development (Galles et al., 2018). Thus, the upregulation of putative ECRs in L3 stages of parasitic nematodes could suggest an analogous role for EC signalling in parasite growth and development at a critical stage in the parasitic lifecycle.

Upregulation of putative ECRs, and a key ECE (*dagl-2*) associated with EC ligand biosynthesis, is evident in *Strongyloides* iL3s (see Figure SI 2). In contrast, the ECEs responsible for EC ligand degradation (*abhd-12* and *faah-1-4*) are downregulated at the iL3 stage (see Figure SI 2). The opposite expression profile is noted in the adult life stage (free-living and parasitic females) of *Strongyloides* spp. where EC-degradation enzymes are upregulated and putative ECRs and *dagl-2* are downregulated (see Figure SI 2). These data suggest that higher levels of EC ligands may exist in the iL3 stage of *S. ratti* and *S. stercoralis* and is consistent with the elevated production of EC-ligands by *N. brasiliensis* iL3s (Batugedara et al., 2018). Together these data indicate that the ECS system may be involved in processes linked to host infection. Parasitic nematodes exploit numerous sensory cues and mechanisms in order to find their host (Gang et al., 2020), thus the upregulation of EC-effectors in iL3s may also implicate EC signalling in sensory perception, host-seeking, and establishment of host infection. These data will direct future functional genomics studies around the role of EC signalling in host finding and infection in tractable parasitic nematodes.

Expression profiling of EC-effectors in sex-specific transcriptome data reveal differential expression patterns in male and female nematodes of several species (see Figure SI 2). In *T. circumcincta* EC- ligand degradation enzymes are broadly downregulated in adult males, and upregulated in adult females (Figure SI 2). Conversely, *O. volvulus* exhibits upregulation of all pathway components in adult males, and downregulation in adult females (Figure SI 2). Sex-specific expression of EC-effectors is common in mammalian species, where they exhibit alternative actions on neuropsychiatric processes and reproductive events (Karasu et al., 2011, Huang and Woolley, 2012, Viveros et al., 2012). In addition the ECS pathway has been implicated in mammalian fertility regulation (Kolodny et al., 1974, Nielsen et al., 2019, Schuel, 2006, El-Talatini et al., 2009, Maccarrone, 2009) and in invertebrate reproduction (Schuel et al., 1991, Schuel et al., 1994). Interrogation of expression at the tissue level is challenging in nematodes where data sets are limited to species which are readily amenable to dissection, for example *Ascaris suum* and *Dirofilaria immitis*. Whilst it would appear that EC-effectors are upregulated in reproductive tissues (e.g. in *A. suum*), further analysis across more tissue types and species is required before meaningful comparisons can be made (data not shown). While the role of EC signalling in the regulation of vertebrate and invertebrate reproduction has been documented, ECS system function in nematode reproduction is yet to be determined. Indeed, enhancing the ability to generate life- and tissue-specific data from key parasitic nematodes will inform functional biology.

## CONCLUSIONS

*In silico* approaches and the proliferation of nematode omics resources provide a valuable opportunity to identify putative novel anthelmintic drug targets for the control of parasite disease. This study focuses on the characterisation of the nematode ECS pathway, driven by its putative biological importance and therapeutic appeal (McPartland and Glass, 2003, Lehtonen et al., 2008, Lucanic et al., 2011, Oakes et al., 2017, Pastuhov et al., 2012). Here we: (i) provide a comprehensive pan-phylum overview of EC-effector complements in nematodes, that represent divergent clades and lifestyles; (ii) unravel the complexity of the nematode ECS to identify putative species- and lifestyle-specific EC-pathways and drug target selectivity; (iii) reveal life stage-, and sex-specific EC-effector expression patterns in relevant parasite species. These data will direct the selection of novel ECS pathway targets for functional validation efforts in parasitic nematodes to inform biology and anthelmintic drug discovery pipelines.

## DATA AVAILABILITY STATEMENT

The original contributions presented in the study are included in the article/Supplementary Material. Further inquiries can be directed to the corresponding author.

## AUTHOR CONTRIBUTIONS

LA, AM, AGM, and NM designed the research. BC, DM and LC, performed the research. BC, DM and LC, analysed the data with assistance from CM and PM. LA, AM, BC, AGM and NM wrote the manuscript. All authors contributed to the article and approved the submitted version.

## ACKNOWLEDGEMENTS

The authors wish to thank WormBase ParaSite for helpful assistance with transcriptome resources.

## ABBREVIATIONS

2-AG: 2-arachidonoylglycerol
ABHD-4: Abhydrolase Domain Containing 4, N-Acyl Phospholipase B
ABHD-5: Abhydrolase Domain Containing 5
ABHD-6: Abhydrolase Domain Containing 6
ABHD-12: Abhydrolase Domain Containing 12, Lysophospholipase
AEA: Anandamide/N-arachidonoylethanolamine
CB1: Cannabinoid Receptor 1
CB2: Cannabinoid Receptor 2
DAGL: Diacylglycerol lipase
EC: Endocannabinoid
EC-GPCR: Endocannabinoid G-protein coupled receptor
ECEs: Endocannabinoid Enzymes
ECRs: Endocannabinoid Receptors
ECS: Endocannabinoid signalling
FAAH-1: Fatty Acid Amide Hydrolase 1
FAAH-2: Fatty Acid Amide Hydrolase 2
GDE-1: Glycerophosphodiester phosphodiesterase 1
Glycero-p-AEA: Glycerophosphoanandamide
GPR-55: G-protein coupled receptor 55
HMM: Hidden Markov Model
Lyso-NAPE: N-acyl-1-acyl-lyso-PE
Lyso-PLD: Lysophospholipase D
MAGL: Monoacylglycerol-lipase
MSA: Multiple Sequence Alignment
NAPE: N-arachidonyl phosphatidyl ethanol-phospholipase D
NAPE-1: N-acyl phosphatidylethanolamine-specific phospholipase-1
NAPE-2: N-acyl phosphatidylethanolamine-specific phospholipase-2
NAPE-PLD: N-Acyl Phosphatidyl Ethanolamine specific phospholipase D
NHR-49: Nuclear Hormone Receptor-49
NPR-9: Neuropeptide Receptor-9
NPR-19: Neuropeptide Receptor-19
NPR-32: Neuropeptide Receptor-32
OCR-2: Osin-9 and Capsacin receptor-related 2
OCTR-1: Octopamine Receptor-1
PLA-2: Phospholipase A2
SER-4: Serotonin/Octopamine receptor family-4
TRPV: Transient receptor potential vanilloid channel

## SUPPLEMENTARY DATA

**File SI 1.** *Caenorhabditis elegans* EC-effector gene IDs.

**File SI 2.** Nematode genome and transcriptome accession numbers and citations.

**File SI 3.** Nematode HMM/BLAST hit gene IDs.

**Table SI 1.** Table of EC-effectors included in study.

**Table SI 2.** EC-effector motifs and citations.

**Figure SI 1. Maximum likelihood phylogeny of:** (A) 98 nematode ABHD-12 homologs. *Homo sapien* ABHD-12 [Q8N2K0 (ABD12_HUMAN)] *H. sapien* MAGL [Q99685(MGLL_HUMAN)], *Caenorhabditis elegans* FAAH-*1-4* [WBGene00015047, WBGene00015048, WBGene00019068, WBGene00013232] and *C. elegans* AHO-3 [WBGene00045192; alpha/beta hydrolase containing protein] also included; (B) 85 nematode NPR-19 homologs. *Homo sapien* CB1 and CB2 [P21554 (CNR1_HUMAN), P34972 (CNR2_HUMAN)], *H. sapien* GPR-55 [Q9Y2T6 (GPR55_HUMAN)] and several *C. elegans* biogenic amine receptors (serotonin [SER-1,-3,-6; WBGene00004776, WBGene00004778, WBGene00021897] dopamine [DOP-1-3; WBGene00001052, WBGene00001053, WBGene00020506] and tyramine [TYRA-2 and -3; WBGene00017157, WBGene00006475]) also included. (C) 89 nematode NPR-9 homologs. *Homo sapien* CB1 and CB2 [P21554 (CNR1_HUMAN), P34972 (CNR2_HUMAN)], *H. sapien* GPR-55 [Q9Y2T6 (GPR55_HUMAN)] and several *C. elegans* biogenic amine receptors (serotonin [SER-1,-3,-6; WBGene00004776, WBGene00004778, WBGene00021897] dopamine [DOP-1-3; WBGene00001052, WBGene00001053, WBGene00020506] and tyramine [TYRA-2 and -3; WBGene00017157, WBGene00006475]) also included. (D) 97 nematode NPR-32 homologs. *Homo sapien* CB1 and CB2 [P21554 (CNR1_HUMAN), P34972 (CNR2_HUMAN)], *H. sapien* GPR-55 [Q9Y2T6 (GPR55_HUMAN)] and several *C. elegans* biogenic amine receptors (serotonin [SER-1,-3,-6; WBGene00004776, WBGene00004778, WBGene00021897] dopamine [DOP-1-3; WBGene00001052, WBGene00001053, WBGene00020506] and tyramine [TYRA-2 and -3; WBGene00017157, WBGene00006475]) also included. (E) 78 nematode NHR-49 homologs. *Homo sapien* PPARG [P37231 (PPARG_HUMAN)], *H. sapien* PPARD [Q03181 (PPARD_HUMAN)], *H. sapien* PPARA [Q07869 (PPARA_HUMAN)], *C. elegans* NHR-88 [WBGene00003678], *C. elegans* NHR-64 [WBGene00003654] and *C. elegans* NHR-35 [WBGene00003628] also included. (F) 107 nematode OCR-2 homologs. *Homo sapien* TRPV1-3 [Q8NER1 (TRPV1_HUMAN), Q9Y5S1 (TRPV2_HUMAN), Q8NET8 (TRPV3_HUMAN)], alongside *C. elegans* OSM-9 [WBGene00003889], *C. elegans* OCR-1 [WBGene00003838], *C. elegans* OCR-3 [WBGene00003840] and *C. elegans* UNC-44 [WBGene00006780] also shown. (G) 99 nematode OCTR-1 homologs. *Homo sapien* CB1 and CB2 [P21554 (CNR1_HUMAN), P34972 (CNR2_HUMAN)], *H. sapien* GPR-55 [Q9Y2T6 (GPR55_HUMAN)], *H. sapien* ADA2A-C [P08913 (ADA2A_HUMAN). P18089 (ADA2B_HUMAN), P18825 (ADA2C_HUMAN)] and several *C. elegans* biogenic amine receptors (serotonin [SER-1,-3,-6; WBGene00004776, WBGene00004778, WBGene00021897] dopamine [DOP-1-3; WBGene00001052, WBGene00001053, WBGene00020506] and tyramine [TYRA-2 and -3; WBGene00017157, WBGene00006475]) also shown. (H) 100 nematode SER-4 homologs. *Homo sapien* CB1 and CB2 [P21554 (CNR1_HUMAN), P34972 (CNR2_HUMAN)], *H. sapien* GPR-55 [Q9Y2T6 (GPR55_HUMAN) and several *C. elegans* biogenic amine receptors (serotonin [SER-1,-3,-6; WBGene00004776, WBGene00004778, WBGene00021897] dopamine [DOP-1-3; WBGene00001052, WBGene00001053, WBGene00020506] and tyramine [TYRA-2 and -3; WBGene00017157, WBGene00006475]) also shown. Outer colours represent nematode clade and circles represent branch support values. Tree generated from an alignment trimmed to include functional domains. Branch supports indicate statistical support from approximate likelihood ratio test (aLRT).

**Figure SI 2. Life-stage and sex-specific expression profiles of EC-effectors.** (A) Ancylostoma ceylanicum, (B) Teladorsagia circumcincta, (C) Dictyocaulus viviparus, (D) Haemonchus contortus, (E) Strongyloides ratti, (F) Strongyloides stercoralis, (G) Trichuris muris and (H) Onchocerca volvulus. Expression heatmaps generated from log2TPM values of all EC-effector transcripts. Average Clustering Method & Pearson’s Distance Measurement Method employed. Life stage and sex-specific data are arranged in columns, rows indicate individual EC-effector transcripts as denoted by effector abbreviation/gene ID. Coloured circles represent EC-receptors (orange), EC biosynthesis enzymes (green) and EC degradation enzymes (purple). [Life stages include; L1, L2, activated L3 (L3A), not activated L3 (L3NA), untreated L3 (L3UT), adult female (AF), adult male (AM) hypobiotic larvae (Lhyp), mixed L1 & L2 (L1+L2), pre-adult L5 female (L5AF), pre-adult L5 male (L5AM), pre-adult L5 mixed gender (L5 Mixed), Adult L5 (L5A), infective larvae (iL3), free-living female (FL Female), tissue migrating L3 (L3+), parasitic female (P Female), post-free living L1 (PFLL1), post-parasitic L1 (PPL1), post-parasitic L3 (PPL3)].

