## Supplementary material for "Pan-phylum *In Silico* Analyses of Nematode Endocannabinoid Signalling Systems Highlight Novel Opportunities for Parasite Drug Target Discovery": Table SI 1

| **EC Effector in *C. elegans* (Proposed Human Ortholog)** | **Proposed Role in nematodes (predominantly derived from *C. elegans*)** | **Citations** |
| --- | --- | --- |
| NPR-9 [G-protein Coupled Receptor 55 (GPR55)] | - Initiation and fine-tuning of backwards locomotion and reversals - Negative regulation of the immune response - Roles in roaming and foraging behaviours | (Bendena et al., 2008, Campbell et al., 2016, Yu et al., 2018) |
| NPR-19 [Cannabinoid Receptor 1 (CB1), GPR55] | - Roles in nociception, locomotion and feeding - Inhibition of axon regeneration | (Pastuhov et al., 2016, Oakes et al., 2017) |
| NPR-32 (CB1) | - Inhibition of axon regeneration | (Pastuhov et al., 2016) |
| NHR-49 [Peroxisome proliferator-activated receptor gamma (PPARY)] | - Control of consumption and balance of fat - Oxidation and desaturation of fatty acids | (Atherton et al., 2008) |
| OCR-2 [Transient Receptor Potential Cation Channel Subfamily V Member 1 [TRPV1)] | - Implicated in larval starvation survival and adult lifespan - Chemosensory functions - Inhibition of aversive behaviour | (Jose et al., 2007, Lee and Ashrafi, 2008) |
| SER-4 [5-Hydroxytryptamine 1A (HTR1A)] | - Roles in nociception and locomotion | (Oakes et al., 2017) |
| OCTR-1 [Alpha-2A adrenergic receptor (ADRA2A)] | - Regulation of innate immunity via suppression of translation - Nociception and locomotion | (Liu et al., 2016, Oakes et al., 2017) |
| NAPE-1/NAPE-2 (N-acyl-phosphatidylethanolamine-hydrolysing phospholipase D) | - NAE biosynthesis - Over-expression has a temperature dependent effect on development/life-span | (Harrison et al., 2014) |
| FAAH-1 (Fatty acid amide hydrolase 1) | - NAE degradation - Over-expression resulted in developmental delays | (Harrison et al., 2014) |
| FAAH-2 (Fatty acid amide hydrolase 2) | - NAE degradation | (Harrison et al., 2014) |
| DAGL-2 (Diacylglycerol lipase 2) | - Synthesis of 2-AG - Over-expression extends mean lifespan by up to 13% | (Lin et al., 2014) |
| ABHD-12 (Monoacylglycerol lipase 2) | - Degradation of 2-AG | (Savinainen et al., 2012, Wei et al., 2016) |
