## Supplementary material for "Pan-phylum *In Silico* Analyses of Nematode Endocannabinoid Signalling Systems Highlight Novel Opportunities for Parasite Drug Target Discovery": Table SI 2

| **EC-effector in *C. elegans* (Proposed Human Ortholog)** | **Functional motif(s)** | **Key amino acid residues** | **Proposed role(s) of motifs** | **Proposed role(s) of key amino acid residues** | **Citations** |
| --- | --- | --- | --- | --- | --- |
| NPR-9 (G-protein Coupled Receptor 55 [GPR55]) | DRY  CWXP  S(N)LAXXAD  NPXXY | Y101, P241, F246 | - DRY – highly conserved motif on transmembrane helix (TMH) 3, role in receptor activation and stabilisation of active states and influence on signal transduction - CWXP – putative molecular hinge essential for recognition of ligands - S(N)LAXXAD – roles in stabilisation of interhelical interaction between TMH2-TMH3/TMH4 - NPXXY- critical for receptor activation | - Y101, P241, F246 – conserved amino acids between GPR-55 and NPR-9; linked directly to AEA binding, Y101 acts as a toggle switch | (Ballesteros and Weinstein, 1995, Pei et al., 2008, Shim, 2009, Shim et al., 2011, Stadel et al., 2011, Lingerfelt et al., 2017, Zhang et al., 2018, Kumar et al., 2019) |
| NPR-19 (Cannabinoid Receptor 1 [CB1], GPR55) | DRY  CWXP  S(N)LAXXAD  NPXXY | F189, K192, L193, F379, S383 | - DRY – highly conserved motif on transmembrane helix (TMH) 3, role in receptor activation and stabilisation of active states and influence on signal transduction - CWXP – putative molecular hinge essential for recognition of ligands - S(N)LAXXAD – roles in stabilisation of interhelical interaction between TMH2-TMH3/TMH4 - NPXXY- critical for receptor activation | - F189 – interacts with AEA amide oxygen; mutation in CB1 decreases AEA binding and affinity sixfold - K192 – interacts with AEA amide oxygen - S383 – forms a hydrogen bond with AEA hydroxyl - F189, L193, F379, S383 – included in AEA binding pocket | (Ballesteros and Weinstein, 1995, McAllister et al., 2003, McAllister et al., 2004, Pei et al., 2008, Shim, 2009, Reggio, 2010, Shim et al., 2011, Stadel et al., 2011, Oakes et al., 2017, Zhang et al., 2018) |
| NPR-32 (CB1) | DRY  CWXP  S(N)LAXXAD  NPXXY | N46, D88, K115 | - DRY – highly conserved motif on transmembrane helix (TMH) 3, role in receptor activation and stabilisation of active states and influence on signal transduction - CWXP – putative molecular hinge essential for recognition of ligands - S(N)LAXXAD – roles in stabilisation of interhelical interaction between TMH2-TMH3/TMH4 - NPXXY- critical for receptor activation | - N46, D88, K115 – implicated in AEA binding | (Ballesteros and Weinstein, 1995, Pei et al., 2008, Shim, 2009, Shim et al., 2011, Stadel et al., 2011, Pastuhov et al., 2016, Zhang et al., 2018) |
| NHR-49 (Peroxisome proliferator-activated receptor gamma [PPARY]) | 9aaTAD motif [positions 495-503] | Q286, S289, H323, H449, Y473 | - 9aaTAD motif – activates transcription as a small peptide; conserved across nuclear hormone receptors | - Q286, S289, H323, H449, Y473 – ligands form hydrogen bonds with these residues in a hydrophilic pocket implicated in agonist binding | (Sheu et al., 2005, Atherton et al., 2008, Piskacek et al., 2019) |
| OCR-2 (Transient Receptor Potential Cation Channel Subfamily V Member 1 [TRPV1]) | CRAC motif [YYTR; positions 553-557] | R491, Y511, S512, T550, R557, E637, D647, E649 | - CRAC motif [YYTR] – conference of cholesterol sensitivity | - T550, Y511 – conference of vanilloid sensitivity and ligand binding - R491, Y511, S512 – molecular determinants and role in ligand binding - R557 – role in volted gate channel gating - E637, D647, E649 – alter receptor sensitivity to divalent cations | (Gavva et al., 2004, Jose et al., 2007, Fernandez-Ballester and Ferrer-Montiel, 2008, Picazo-Juarez et al., 2011) |
| SER-4 (5-Hydroxytryptamine 1A [HTR1A]) | DRY  CWXP  S(N)LAXXAD  NPXXY | I113, D116, V117, I189, F361, F362, Y390 | - DRY – highly conserved motif on transmembrane helix (TMH) 3, role in receptor activation and stabilisation of active states and influence on signal transduction - CWXP – putative molecular hinge essential for recognition of ligands - S(N)LAXXAD – roles in stabilisation of interhelical interaction between TMH2-TMH3/TMH4 - NPXXY- critical for receptor activation | - I113, V117, F361, F362 – hydrophobic interactions with ligands via nonpolar side chain - D116 – formation of salt bridge with ligand - I189 – forms hydrogen bonds with indole ring moiety - Y390 – forms hydrogen bond with centre of ligands | (Ballesteros and Weinstein, 1995, Pei et al., 2008, Shim, 2009, Stadel et al., 2011, Oakes et al., 2017, Zheng et al., 2017, Zhang et al., 2018) |
| OCTR-1 (Alpha-2A adrenergic receptor [ADRA2A]) | DRY  CWXP | D94, D128, D145, S214, S219, F423, F427 | - DRY – role in receptor activation and stabilisation of active states and influence on signal transduction - CWXP – putative molecular hinge essential for recognition of ligands | - D94, D128, D145, S214, S219, F427 – Role in ligand binding. Mutagenesis of these residues results in lower affinity for receptor agonists - F423 – in binding pocket | (Suryanarayana et al., 1991, Wang et al., 1991, Ballesteros and Weinstein, 1995, Pei et al., 2008, Oakes et al., 2017) |
| NAPE-1/NAPE-2 (N-acyl-phosphatidylethanolamine-hydrolysing phospholipase D) | HX(E/H)XD(C/R/S/H)X50–70HX15–30(C/S/D)X30–70H signature sequence | H253, D284, Q320, H321, H343 | - HX(E/H)XD(C/R/S/H)X50–70HX15–30(C/S/D)X30–70H signature sequence – role in zinc binding and hydrolysis | - H253, D284, Q320, H321, H343 – key involvement in catalytic processes | (Okamoto et al., 2004, Harrison et al., 2014) |
| FAAH-1 (Fatty acid amide hydrolase 1) | Amidase signature domain | K142, M191, S217, S241 | - Amidase signature domain – catalytic motif | - K142, S217, S241 – catalytic triad - M191 – inside active site | (Lucanic et al., 2011, Haq and Kilaru, 2020) |
| FAAH-2 (Fatty acid amide hydrolase 2) | Amidase signature domain | K131, C180, S206, S230 | - Amidase signature domain – catalytic motif | - K131, S206, S230 – catalytic triad - C180 – inside active site | (Lucanic et al., 2011, Sirrs et al., 2015, Haq and Kilaru, 2020) |
| DAGL-2 (Diacylglycerol lipase 2) | Serine lipase domain  PPXXF | S472, D524 | - Serine lipase domain – catalytic domain - PPXXF – consensus motif for binding coiled-coil domain of Homer proteins | - S472, D524 – inside active site | (Reisenberg et al., 2012) |
| ABHD-12 (Monoacylglycerol lipase 2) | Alpha/beta hydrolase  Domain [consists of  lipase motif (GTSMG)] | S122,S148, D230, D278, H269, H306 | - Lipase motif (GTSMG) – catalytic domain | - S122, D230, H269 – active site identified from mutagenesis studies - S148, D278, H306 – catalytic triad | (Karlsson et al., 1997, Savinainen et al., 2012) |
