## Supplementary figures and images for "Pan-phylum *In Silico* Analyses of Nematode Endocannabinoid Signalling Systems Highlight Novel Opportunities for Parasite Drug Target Discovery"

### Figure SI 1

(A)

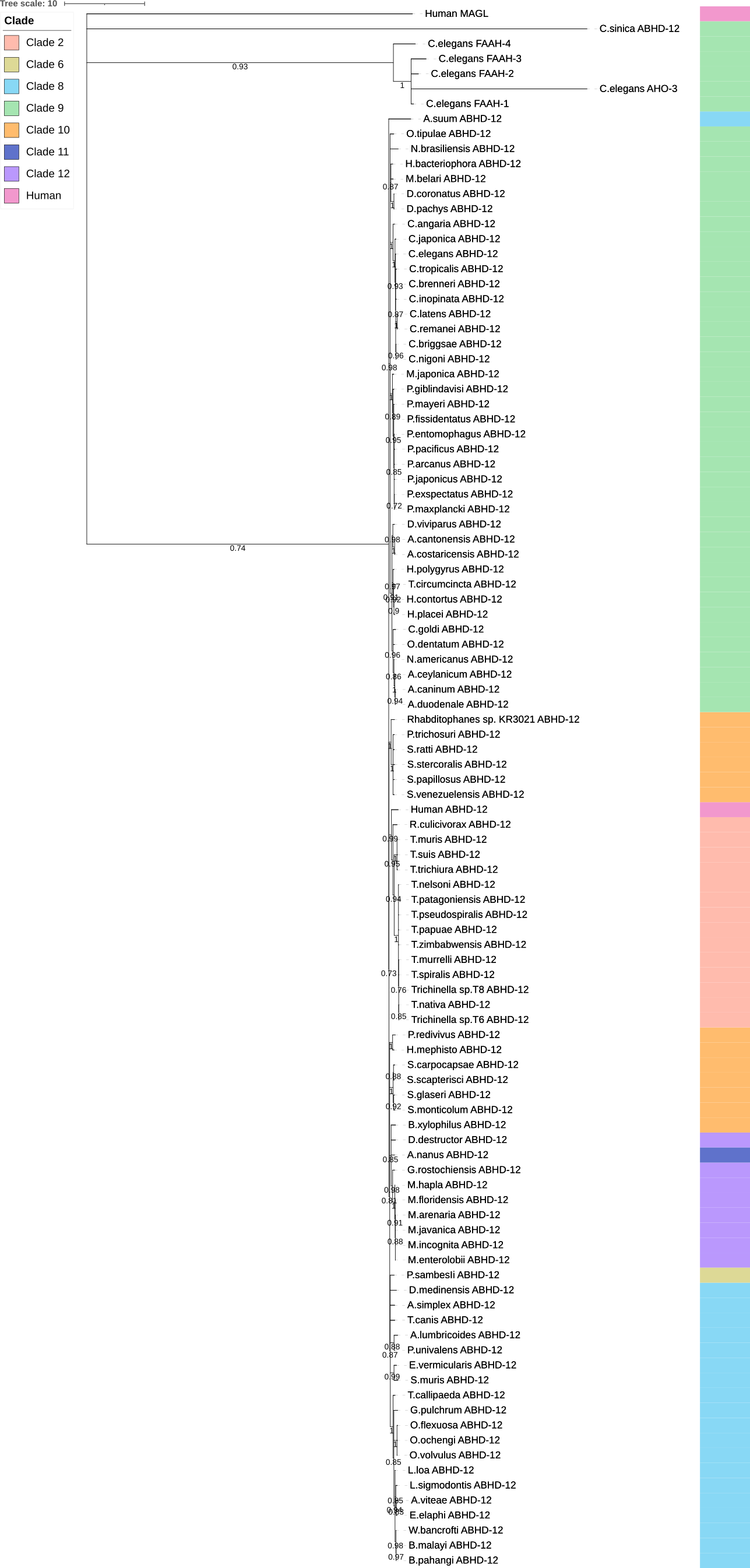

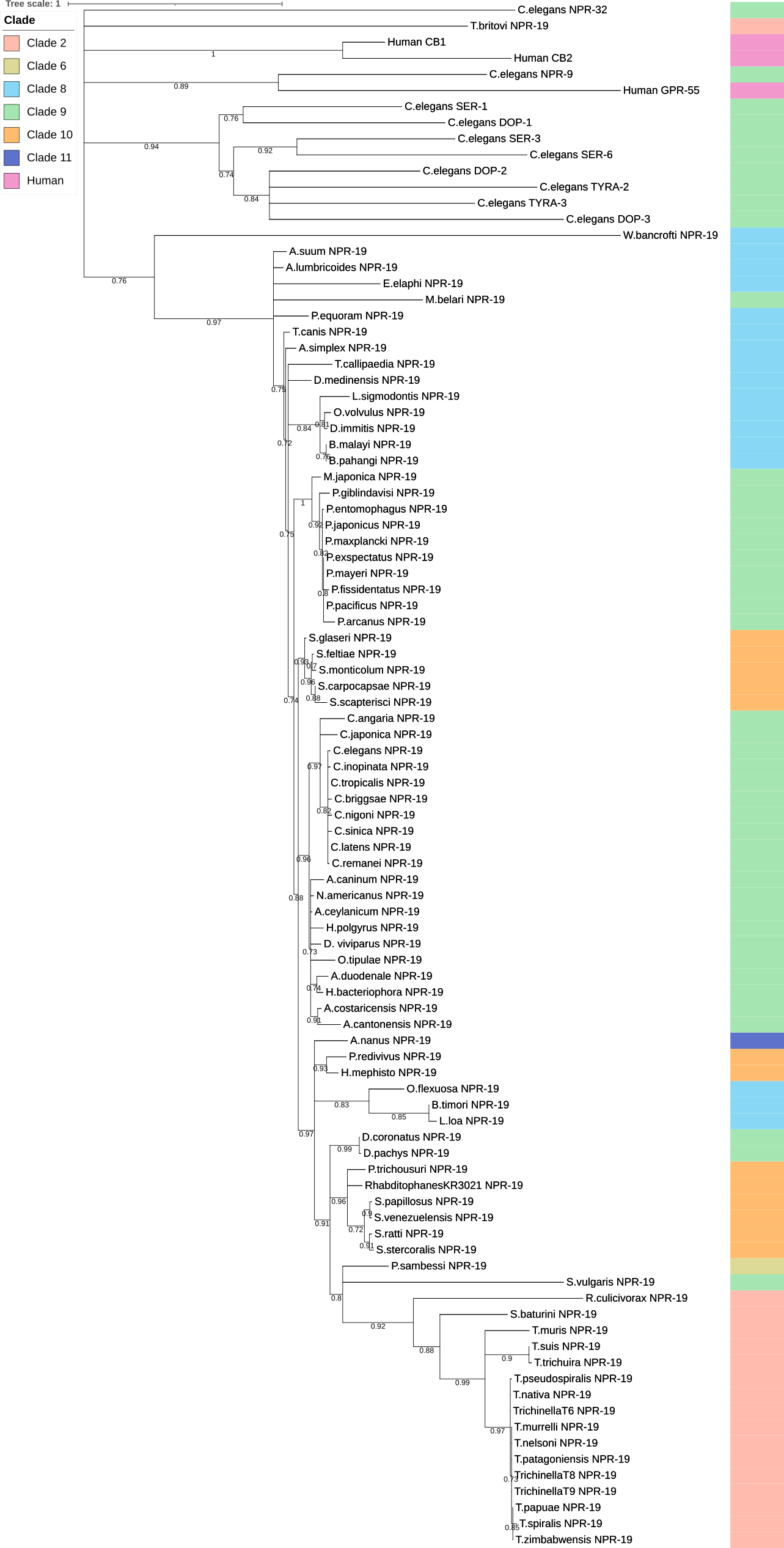

(B)

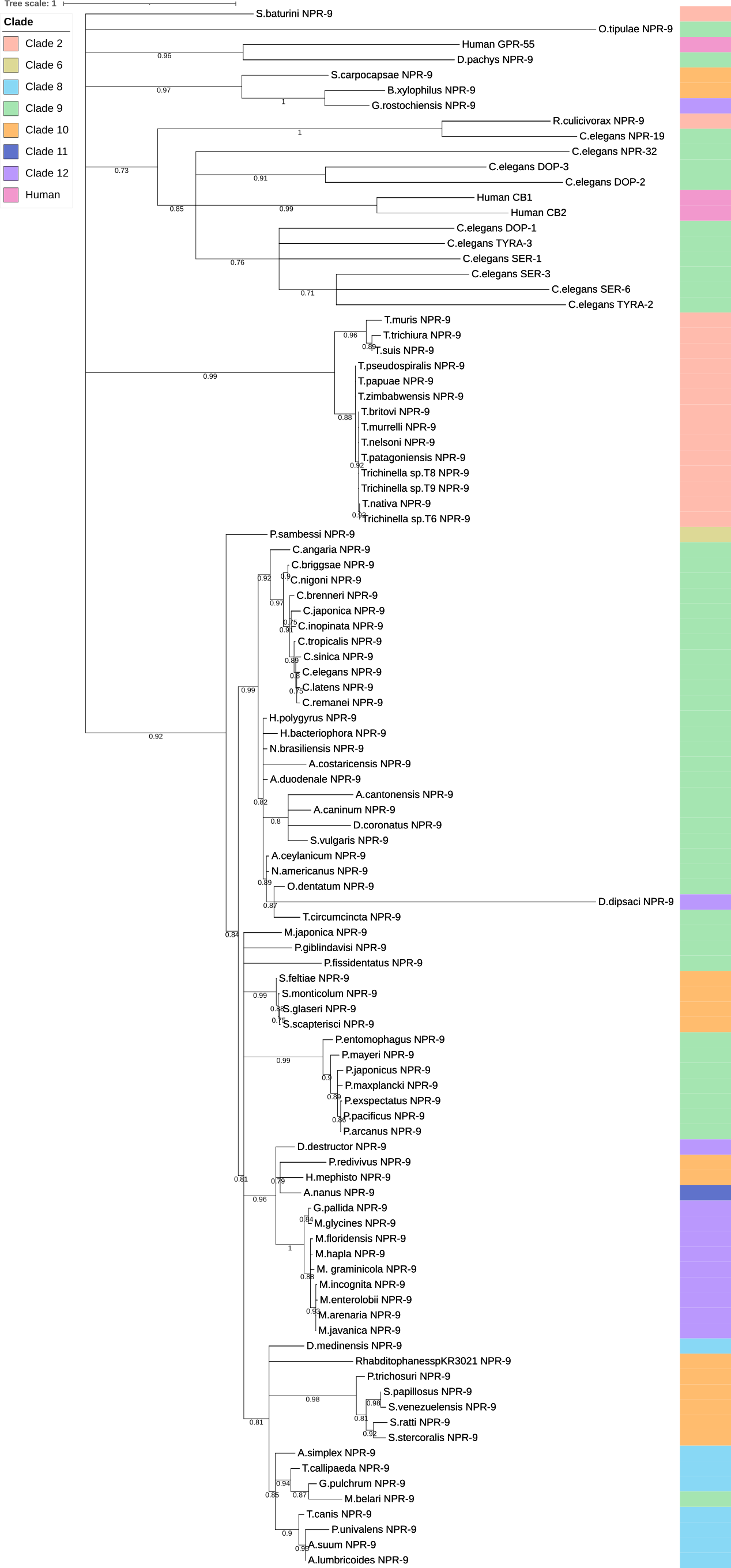

(C)

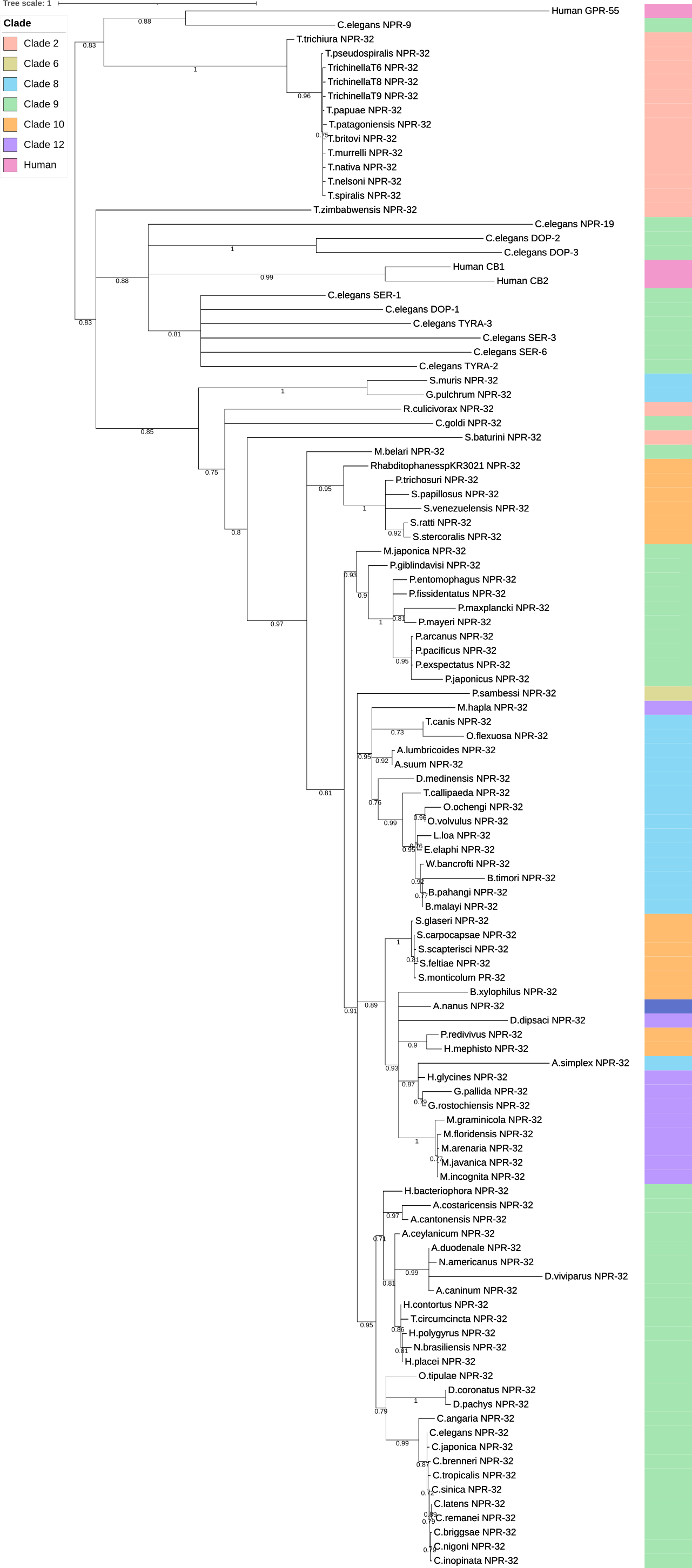

(D)

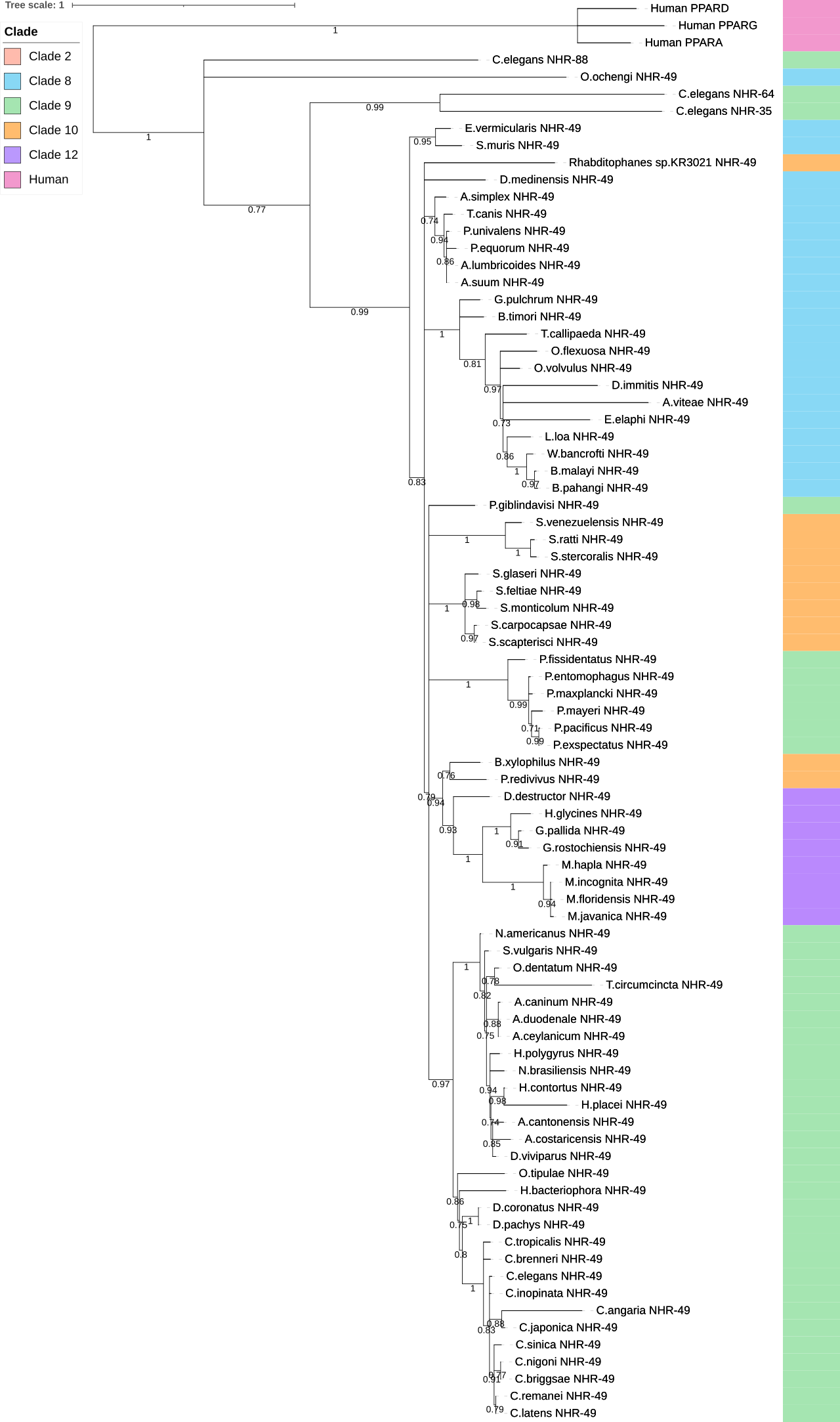

(E)

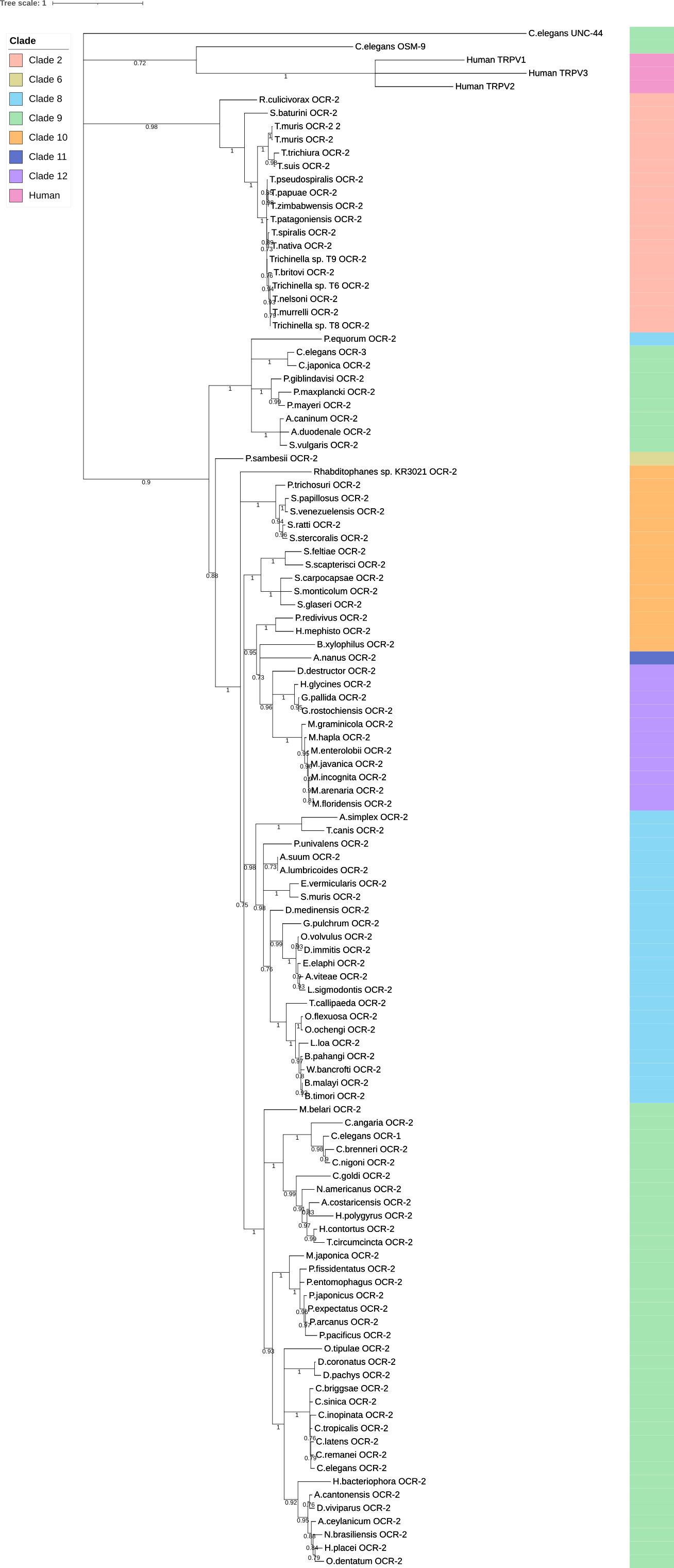

(F)

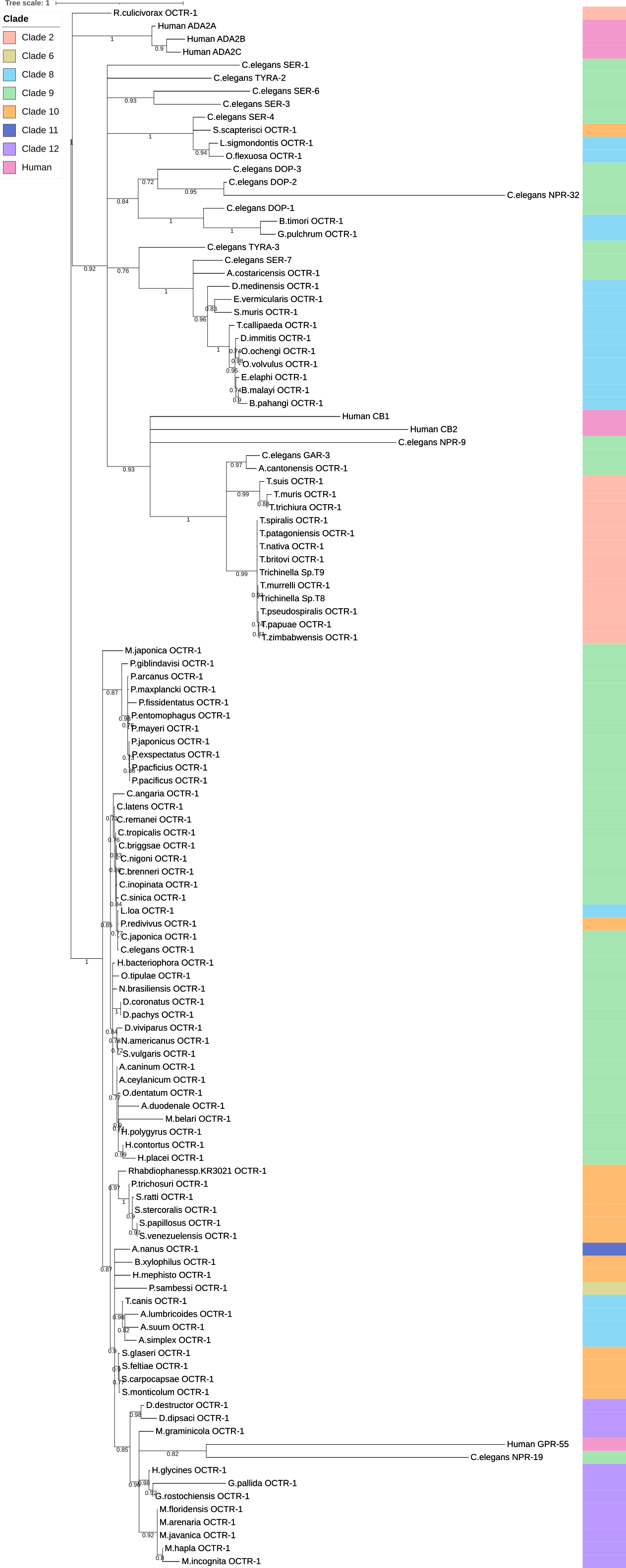

(G)

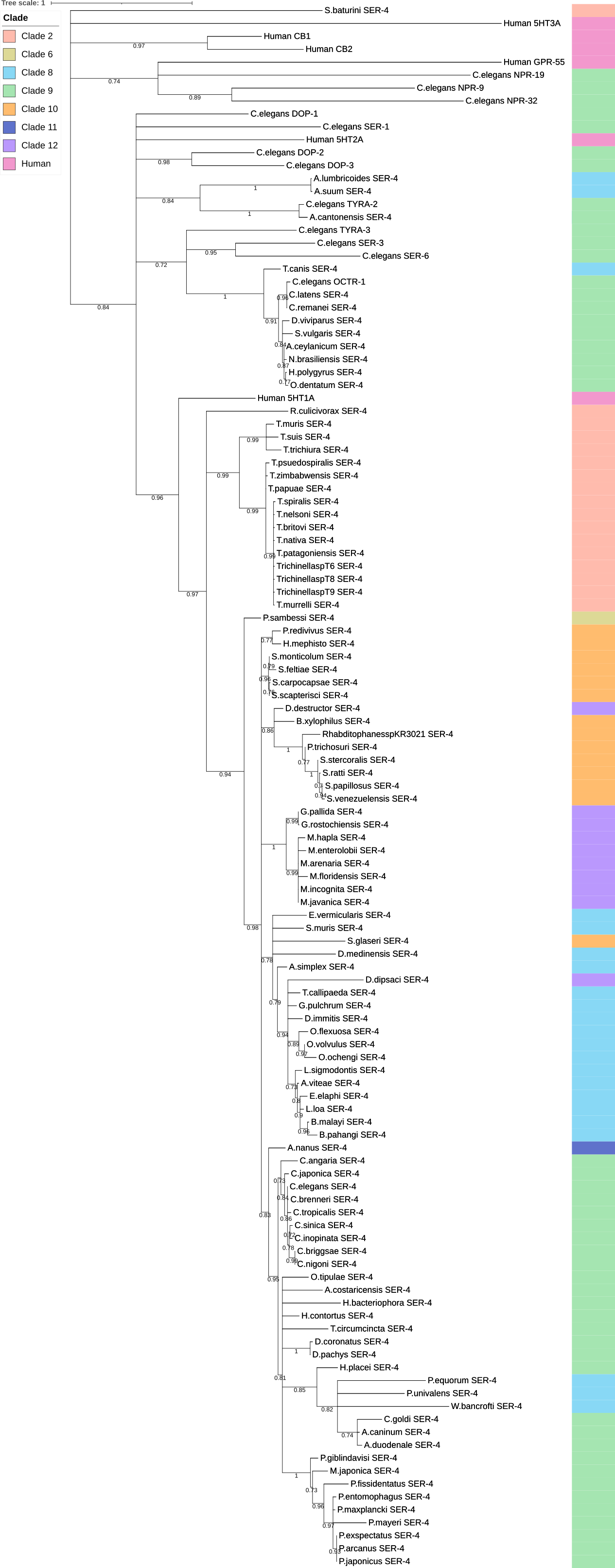

(H)

### Figure SI 2

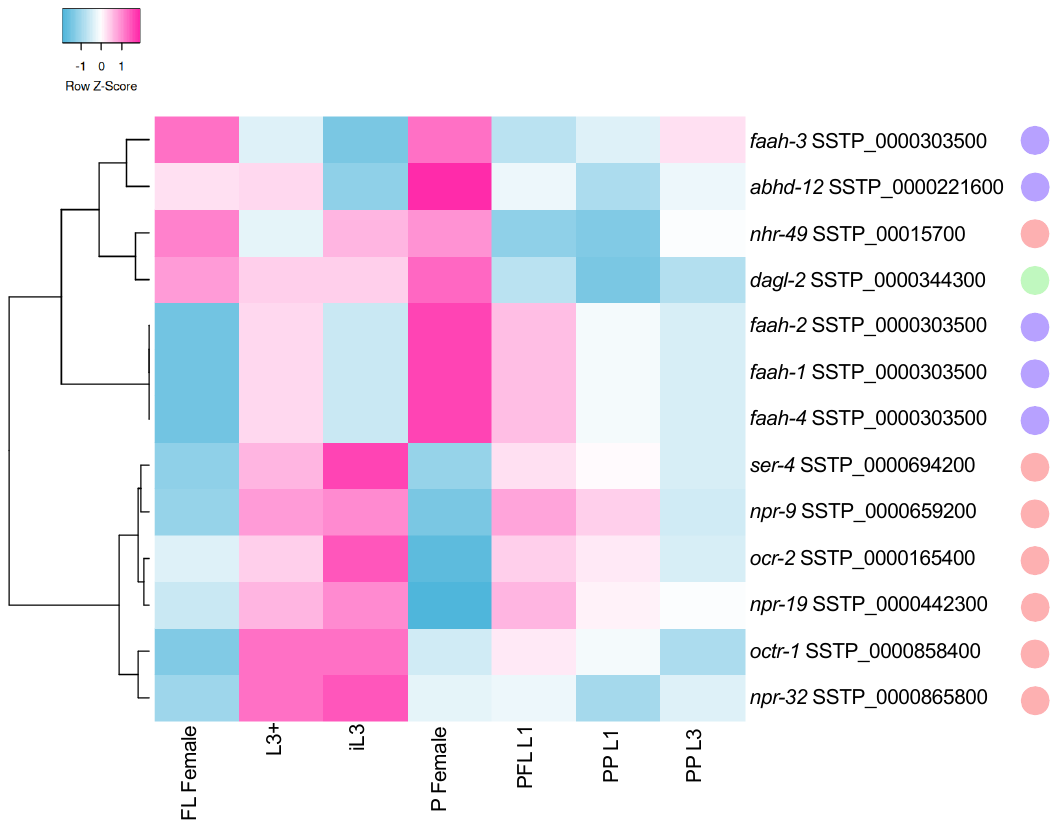

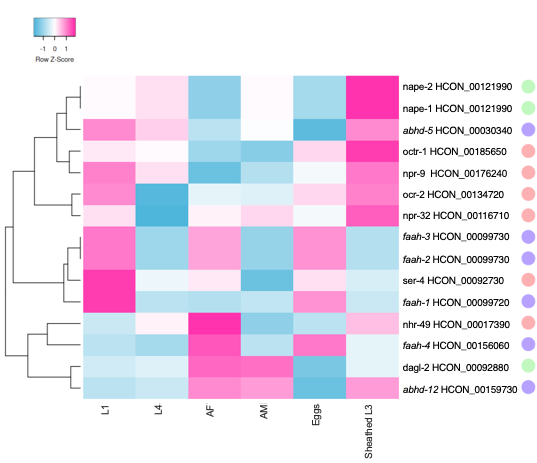

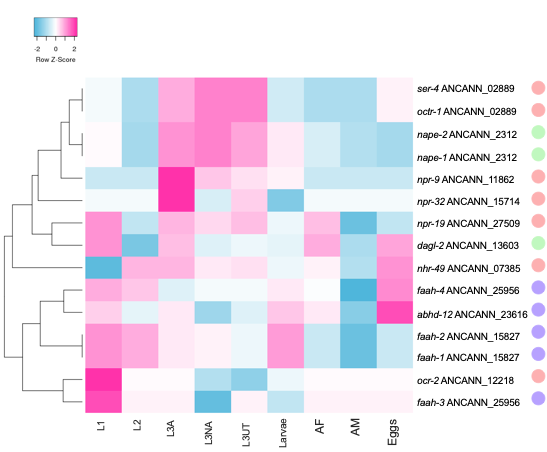

**A**

**B**

**C**

**D**

**E**

**F**

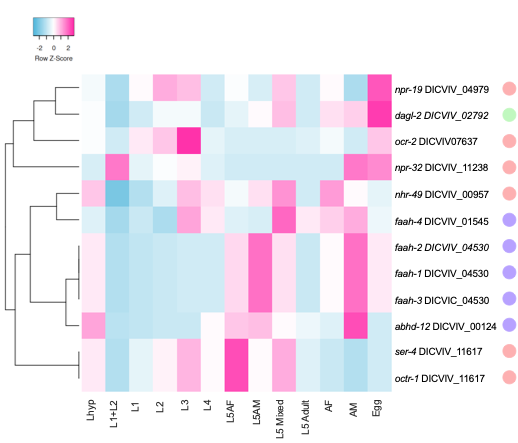

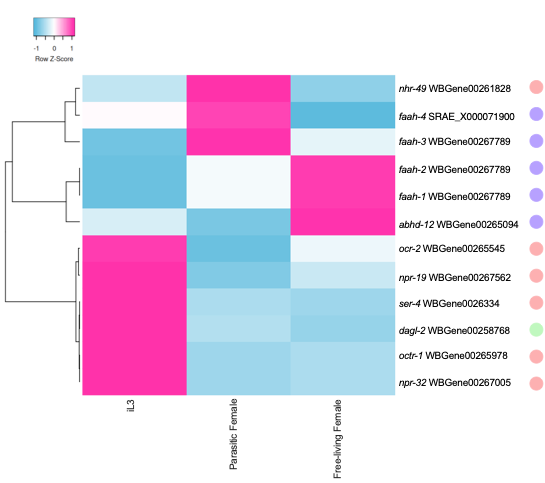

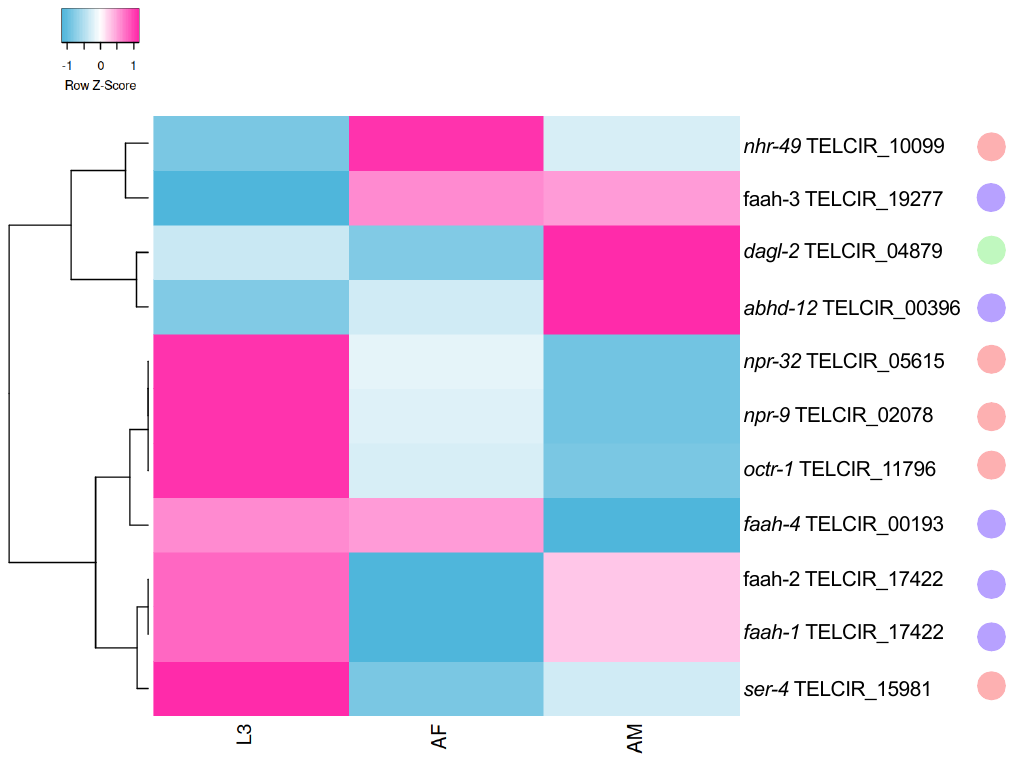

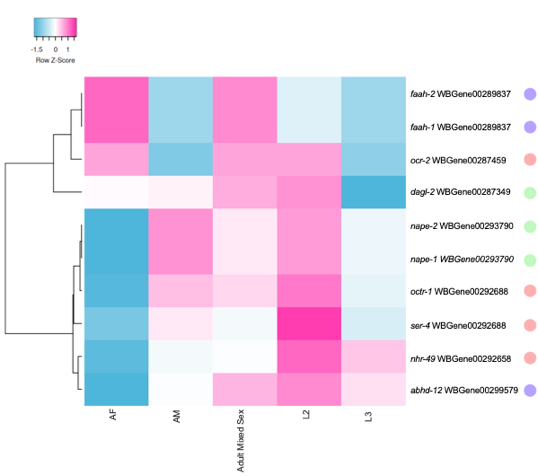

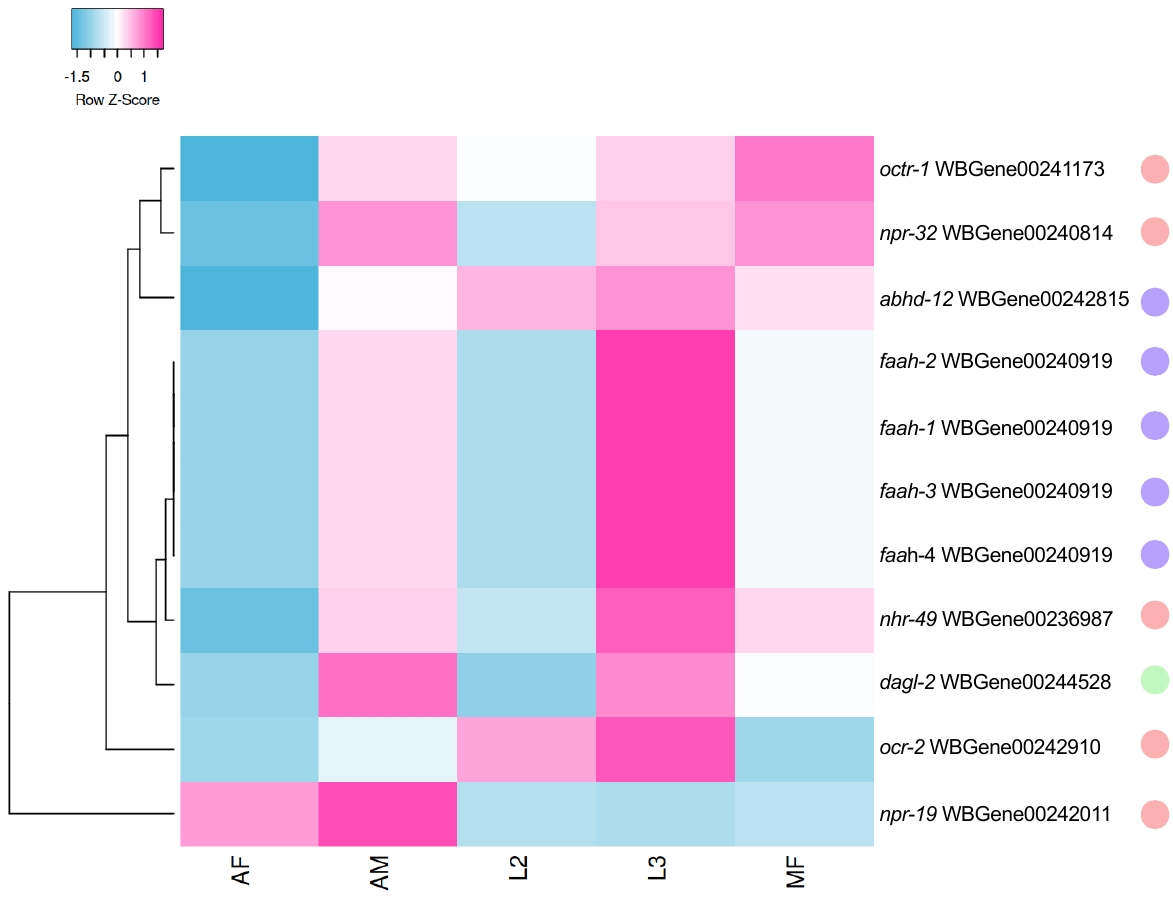

**H**

**G**
